# Behavioral signatures of face perception emerge in deep neural networks optimized for face recognition

**DOI:** 10.1101/2022.11.23.517478

**Authors:** Katharina Dobs, Joanne Yuan, Julio Martinez, Nancy Kanwisher

## Abstract

Human face recognition is highly accurate, and exhibits a number of distinctive and well documented behavioral “signatures” such as the use of a characteristic representational space, the disproportionate performance cost when stimuli are presented upside down, and the drop in accuracy for faces from races the participant is less familiar with. These and other phenomena have long been taken as evidence that face recognition is “special”. But why does human face perception exhibit these properties in the first place? Here we use deep convolutional neural networks (CNNs) to test the hypothesis that all of these signatures of human face perception result from optimization for the task of face recognition. Indeed, as predicted by this hypothesis, these phenomena are all found in CNNs trained on face recognition, but not in CNNs trained on object recognition, even when additionally trained to detect faces while matching the amount of face experience. To test whether these signatures are in principle specific to faces, we optimized a CNN on car discrimination and tested it on upright and inverted car images. As for face perception, the car-trained network showed a drop in performance for inverted versus upright cars. Similarly, CNNs trained only on inverted faces produce an inverted inversion effect. These findings show that the behavioral signatures of human face perception reflect and are well explained as the result of optimization for the task of face recognition, and that the nature of the computations underlying this task may not be so “special” after all.

**Significance Statement:** For decades, cognitive scientists have collected and characterized behavioral signatures of face recognition. Here we move beyond the mere curation of behavioral phenomena to asking why the human face system works the way it does. We find that many classic signatures of human face perception emerge spontaneously in CNNs trained on face discrimination, but not in CNNs trained on object classification (or on both object classification and face detection), suggesting that these long-documented properties of the human face perception system reflect optimizations for face recognition per se, not by-products of a generic visual categorization system. This work further illustrates how CNN models can be synergistically linked to classic behavioral findings in vision research, thereby providing psychological insights into human perception.

## Introduction

For over fifty years, cognitive psychologists have documented the many ways that face recognition is “special” (1, 2). Face recognition performance drops disproportionately for inverted faces (i.e., face inversion effect) (3), is higher for faces of familiar than unfamiliar races (i.e., other-race effect) (4), and makes use of a characteristic “face space” (5). These and other behavioral signatures of the face system have been collected and curated as evidence that qualitatively distinct mechanisms are engaged in the recognition of faces compared to other objects. But largely missing from this long-standing literature is the question of *why* the human face recognition system might have these particular properties. Ideal observer methods have long been used to test whether specific behaviorally-observed phenomena reflect optimized solutions to simple perceptual tasks, but this method is not well suited for complex real-world tasks (6) like face recognition. Recently, however, task-optimized deep neural networks are providing new traction on this classic question (7). In particular, if a specific human behavioral phenomenon is the expected result of optimization for a given task (whether through evolution or individual experience), then we should observe a similar phenomenon in a deep neural network optimized for that same task. Here, we use this logic to test the hypothesis that the classic behavioral signatures of face perception result specifically from optimization for the task of discriminating one face from another, by testing the prediction that these signatures will be found in convolutional neural networks (CNNs) trained on face recognition, but not in CNNs trained on object categorization or face detection, even when their overall face experience is matched.

Reason to suspect that training on faces may be important for CNNs to capture human face perception behavior comes from the ample evidence that face-trained networks perform well on face recognition tasks (8–10). But even if face experience is necessary, it could still be that training on face detection alone (without fine-grained face recognition) is sufficient for specific phenomena in human face perception to emerge. In contrast, reason to suspect that face experience may not be necessary to capture human behavior comes from previous findings that the features learned by CNNs optimized for object recognition are highly predictive of human perceptual similarity (11) and broadly useful for many tasks beyond visual object categorization (12–14). Further, object-trained networks are currently the best model of face-specific neural responses in the primate brain (15, 16), and even appear to contain units selectively responsive to faces (17, 18). A third possibility is that none of the above training regimes might be able to capture all classic signatures of human face perception, and something else might be required, such as a face-specific inductive bias (19, 20) or a higher-level semantic processing of faces (21), to capture human behavioral signatures of face processing. Finally, these hypotheses are not mutually exclusive, and it is possible that different signatures of human face processing may result from optimization for different tasks.

Here, we tested humans in five different experiments on tasks that measure performance on real-world face recognition, “face space”, and two of the classic signatures of human face perception: the face inversion effect and other-race effect (see Table S1-S2 for details). These behavioral face perception signatures were then directly compared to multiple CNNs based on the same architecture but optimized for different tasks (see Table S3-S4 for details): one network was optimized on fine-grained face recognition, one was trained on object categorization only (without face categories in the output layer), one was trained on object categorization and face detection (assigning all faces to one output category), and one was not trained at all (i.e. the same CNN with random weights). A recent study found that face-trained but not object-trained CNNs approached human face recognition accuracy (22). Here we begin by replicating this phenomenon in a large-scale cohort. We then compared not only overall accuracy, but also the representational similarity space between the networks and humans. Next we asked whether human face signatures emerge only from an optimization for face recognition, or whether an optimization for face detection would suffice. Lastly, we tested whether a classic face signature – the face inversion effect – is specific for faces per se or whether it can, in principle, emerge for other categories in networks optimized for fine-grained discrimination of those categories (23). Critically, although all CNNs necessarily start with a particular architecture and learning rule, none of the networks had any built-in inductive biases – except those introduced by the different objective functions – to produce these specific behavioral signatures.

## Results

### Does humanlike face recognition performance reflect optimization for face recognition in particular?

One of the most basic properties of human face recognition is simply that we are very good at it. Could our excellent accuracy at face recognition result from generic object categorization abilities, or does it reflect optimization for face recognition in particular? In Experiment 1, we measured face recognition performance in people and CNNs using a difficult target-matching task of choosing which of two face images belong to the same identity as a third target image (Fig. 1B, left panel). Target and nontarget faces were all white females between 20-35 years of age, so discrimination of age, gender, or race alone would not suffice for high accuracy on this task. The correctly matching face image differed from the target face image in many low-level features and often in viewpoint, lighting, and facial expression, requiring participants to abstract across these differences to match the identity. Humans (n=1,532) were tested in a large-scale online experiment using Amazon Mechanical Turk.

**Figure 1.**
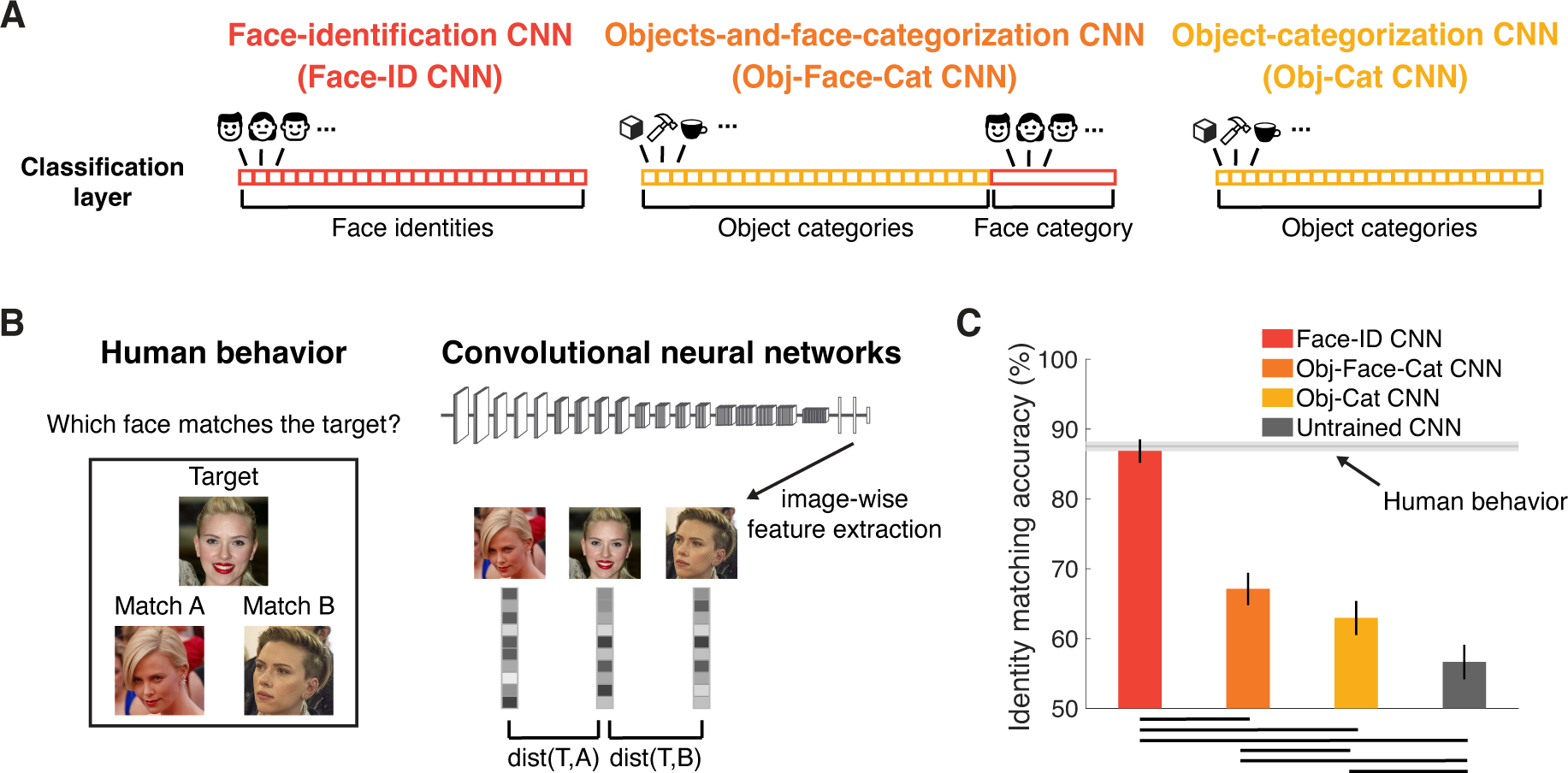
Experiment 1: Only CNNs trained on face recognition achieve human-level accuracy. **(A)** We compared four CNNs with the VGG16 architecture to human behavior: one trained on face identity recognition (Face-ID CNN, red), one on object and face categorization, with all faces used for training the face-ID CNN assigned to one category (Obj-Face-Cat CNN, orange), one trained on object categorization only (Obj-Cat CNN, yellow), and another untrained, randomly initialized CNN (Untrained CNN). **(B)** Human face recognition performance (n=1,532) was measured in a target-matching task on 40 female identities (5 images each) on Amazon Mechanical Turk. To measure performance in CNNs on the same task, we extracted the activation to each of the images in a triplet and computed the correlation distance (1 – Pearson’s *r*) between the target (T) and the two matching images (A and B). The network’s choice was modeled as the minimal distance between the target and each of the matching images (e.g., dist(T,A)). **(C)** Human performance was 87.5% correct (light gray horizontal line; chance level was 50%). Only the face-trained CNN (red) achieved face recognition performance close to humans. Networks trained on object categorization and face detection (orange), or object categorization only (yellow) performed better than untrained CNNs (gray), but did not reach human-level recognition performance. Error bars denote bootstrapped 95% CIs. Black lines on the bottom indicate pairwise significant differences (p<0.05, bootstrap tests, fdr-corrected). Images shown are not examples of the original stimulus set due to copyright; images shown are in public domain and available at https://commons.wikimedia.org.

Four CNNs were tested on the same task. All four were based on the VGG16 architecture (Fig. 1A): one trained to discriminate face identities (Face-ID CNN in red), one trained on object categorization, excluding all animal categories (Obj-Cat CNN in yellow), one trained on object and face categorization (including all the faces and object training images, but assigning all face images to a single category; Obj-Face-Cat CNN in orange), and one untrained, randomly initialized CNN (Untrained CNN in gray). We chose the VGG16 architecture (24) because it provides a good fit to neural visual processing (25), it has been successfully trained for face recognition (26) and is widely used in cognitive neuroscience (see Supplementary Note 2 for similar results from two other commonly-used architectures: Alexnet (27) and ResNet (28, 29)). In CNNs, we extracted activation patterns from the penultimate fully-connected layer (i.e., the decoding stage in a CNN; see Supplementary Note 1 for results from other layers) to the same images and computed the correlation distance (1 – Pearson’s *r*) between the activation patterns of each pair of images (Fig. 1B, right panel). The network’s choice was determined by which of the two matching images had an activation pattern that was closest to the target image. Importantly, none of the networks was trained on the face identities used as test stimuli.

Human participants were able to correctly match the target face in 87.5% of all trials (1C, light-gray horizontal line; chance level was 50%). Although we intentionally chose less well-known faces for this test, it remained possible that our participants recognized some of the individuals, possibly inflating performance. To find out, we asked each participant if one or more of the identities were familiar to them. Indeed, 71% of participants indicated that they were familiar with at least one of the identities, and 10% indicated that they were ‘not sure’. When we ran the same analysis separately on those participants who indicated that they were familiar with at least one identity or ‘not sure’ (n=1,233) versus those that were not familiar with any of the identities (n=299), we found a significant but small drop in performance from 88.3% to 84.5% (p=0, bootstrap test). This finding suggests that any contribution of familiarity with particular faces to the observed performance in this task was small (cf. Fig. 3A for very similar performance on completely unfamiliar faces).

Might this characteristically high human accuracy on a difficult, high-variance face recognition task, result from a system optimized only for generic object categorization, or would specific optimization for faces be required? We found that CNNs trained on object categorization performed far worse (Obj-Cat CNN: 63.0%; Fig. 1C, yellow) than humans, whereas the face-identity trained CNN achieved human-level performance at 86.9% correct (Face-ID CNN; Fig. 1C, red; p=0.44, bootstrap test), consistent with prior studies (8–10, 22, 30). Does the human-level accuracy of the face identity-trained CNN result from an optimization specifically for face identity discrimination, or would a CNN with the same amount of face experience but optimized for coarse face detection also achieve it? We found that the CNN trained to categorize objects and faces (assigning all faces to a single output category) performed significantly better than the CNN trained on object categorization only (Obj-Face-Cat CNN: 67%; Fig. 1C, orange; p<0.01, bootstrap test), but it performed significantly worse than human performance (p=0, bootstrap test). The untrained CNN achieved a performance of 56.7%, which was significantly above chance (Fig. 1C, gray; p=0, bootstrap test) but much lower than human performance (Fig. 1C, light-gray horizontal line) or performance of all other trained networks (all p=0, bootstrap tests).

Does the face-identity trained CNN not just match the overall performance of humans, but also use similar strategies to solve the identity matching task (31, 32)? To address this question, we performed an analysis of the errors being made by humans and CNNs (see Supplementary Note 3 for details). We indeed found that humans performed significantly better on triplets in which the Face-ID CNN was correct (human performance 88.2%) than on triplets for which it was incorrect (human performance: 83.3%; p=0, bootstrap test). This finding suggests that the face-identity trained CNN not only achieves a similar recognition accuracy to humans, but uses a similar strategy to do so.

Thus, we found that CNNs optimized for face recognition were able to achieve face recognition performance comparable to humans (consistent with prior studies (9, 10, 22)), but this was not the case for CNNs trained on object categorization (despite their high usefulness for other tasks (11, 14)) or untrained CNNs. Further, the CNN trained on both object classification and face detection, which had “experienced” as many faces as the CNN optimized for face recognition but was trained to assign all faces to a single face category, performed far worse than the CNN trained on face recognition. Taken together, these findings suggest that humans’ high accuracy at face recognition is not the result of a system optimized for generic object categorization, even with large numbers of faces in the training data, but more likely reflects optimization (through evolution or individual experience) for face recognition in particular.

### Do CNNs represent faces in a similar fashion to humans?

The analyses so far show that CNNs trained on face recognition achieve accuracy levels similar to humans when tested on the same task. But do they achieve this high performance in the same way? To address this question, we assessed the perceived similarity of face images in humans and compared them to CNNs using representational similarity analysis (RSA). Specifically, in Experiment 2, we asked whether the similarity between face representations in CNNs resemble those in humans. Human participants (n=14) performed a multi-arrangement task (33) on images belonging to 16 different identities (5 images each) for a total of 80 face images (Fig. 2A). In this task, participants were asked to place each image in a 2D space that captures similarities in the appearance of faces. Using RSA, we compared the resulting behavioral representational dissimilarity matrices (RDMs) to the RDMs of all three CNNs, obtained by computing the correlation distance between the activation patterns from the penultimate fully-connected layer for the same stimuli (see Supplementary Note 5 for results from other layers and Supplementary Note 6 for similar results from two other architectures: Alexnet and ResNet).

**Figure 2.**
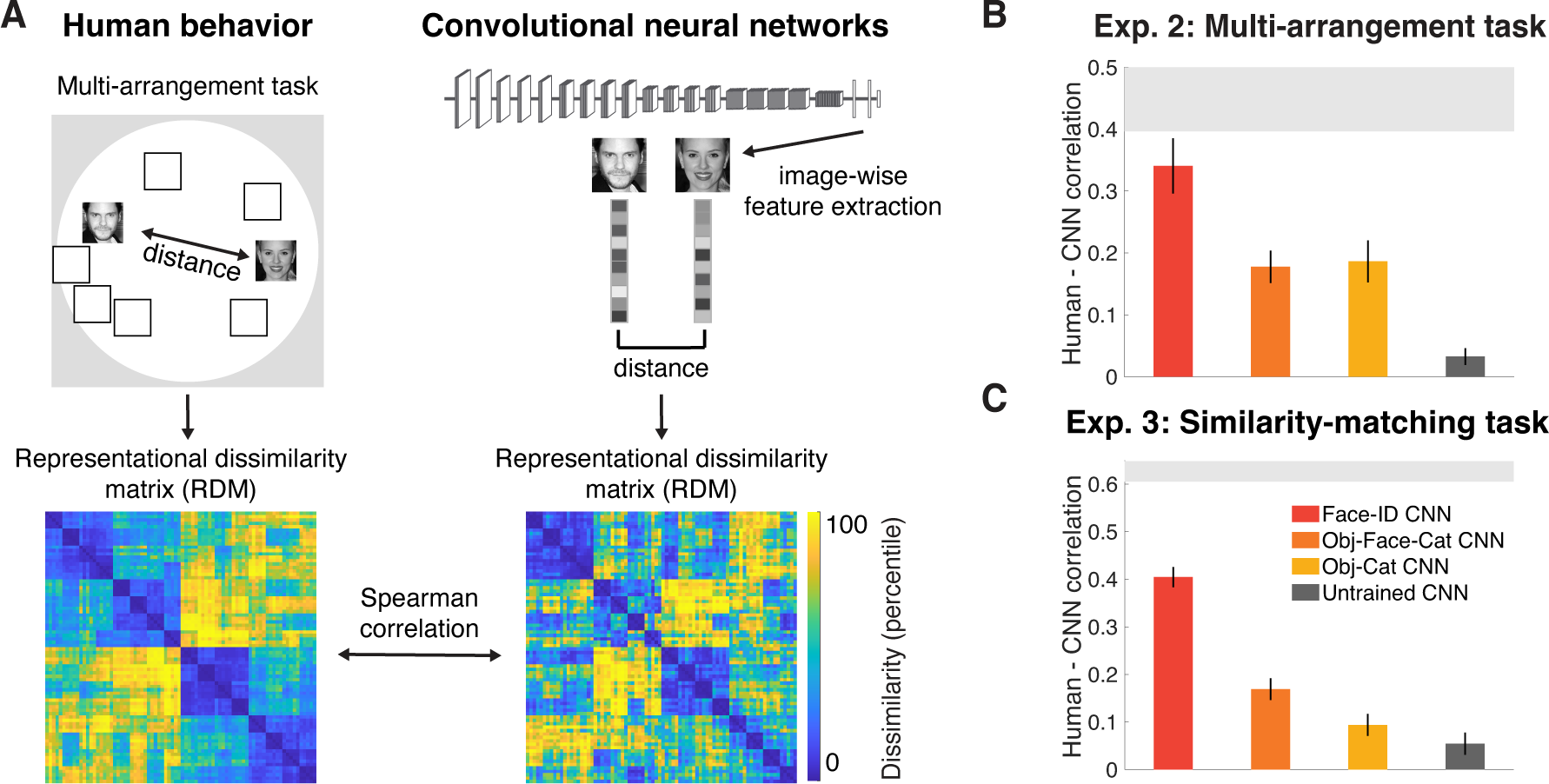
Experiments 2 and 3: Face-trained but not object-trained CNNs match human face behavior. **(A)** To measure human representational similarities of faces in Experiment 2, participants (n=14) performed a multi-arrangement task on 16 face identities (5 images each) resulting in a representational dissimilarity matrix (RDM) for each participant (the average RDM across all participants is shown in the bottom left). The correlation distance (1 – Pearson’s *r*) between activations to each pair of the corresponding images in the penultimate fully-connected layer was used to compute the networks’ RDM (sample RDM is shown for the penultimate layer of the Face-ID CNN in the bottom right, see Supplementary Note 4 for other RDMs). Spearman rank correlation was used to measure the similarity between human behavioral and networks’ RDMs. The 16 face identities were half female, half male and half old, half young. The low dissimilarity clusters (in blue) that can be seen along the diagonal of the RDMs correspond to ‘old female’, ‘young female’, ‘old male’ and ‘young male’ identities (from top left to bottom right), respectively. **(B)** The face-identity trained CNN (Face-ID CNN, red) matched human behavioral representational similarity best (close to noise ceiling; light gray bar). Neither the untrained CNN (Untrained CNN, dark gray) nor the object-categorization trained CNN (Obj-Cat CNN, yellow) or the CNN trained to categorize objects and to detect faces (Obj-Face-Cat CNN, orange) matched human representational similarities well. Error bars represent bootstrapped SEMs across participants. The gray area represents the noise ceiling. **(C)** The results of (B) were replicated in Experiment 3 using a similarity-matching task (see Fig. 1B for the same stimulus presentation methods but different task: identity matching in Experiment 1 but similarity matching here in Experiment 3) on Amazon Mechanical Turk based (n=668) on a distinct dataset of 60 unfamiliar male identities (one image each). The face-trained CNN (Face-ID CNN, red) again matched human behavioral representational similarity best, far outperforming the untrained CNN (gray), the object-categorization trained CNN (Obj-Cat CNN, yellow) and the object-categorization and face-detection trained CNN (Obj-Face-Cat CNN, orange). The corresponding RDMs are shown in Supplementary Note 4. The gray area represents the split-half reliability (mean±2*SD across 50 random splits). Error bars denote bootstrapped 95% CIs across dissimilarity values. Images shown are not examples of the original stimulus set due to copyright restrictions; images shown are in public domain and available at https://commons.wikimedia.org.

For the face-identity trained CNN (Exp. 2; Fig. 2B; Face-ID CNN in red), correlations between the network’s face representations and human behavior were high in the penultimate layer (Spearman’s *r*: 0.34), almost reaching the noise ceiling (i.e., the maximum correlation possible given the consistency across participants; light gray vertical bar). In contrast, the CNN trained object categorization only (Fig. 2B; Obj-Cat CNN in yellow) and the CNN trained on object and face categorization (Fig. 2B; Obj-Face-Cat CNN in orange) represented faces significantly less similarly to humans (Obj-Cat CNN: Spearman’s *r*: 0.19; Obj-Face-Cat CNN: Spearman’s *r*: 0.21; both p=0, bootstrap test). The representational dissimilarities of the untrained CNN (Fig. 2B; Untrained CNN in dark gray) showed a significant but low correlation with human behavior (Spearman’s *r*: 0.03; p=0.02, bootstrap test). Thus, the decoding stage of processing in face-identity trained CNNs, but not CNNs trained on object categorization, face detection or untrained CNNs, match human behavior well, indicating that faces are similarly represented in human behavior and face-identity trained CNNs.

The previous dataset contained multiple images of the same identity, thus human participants might have been biased to simply place images of the same identity together, without taking into account fine-grained details within or between identities. Do these results generalize to other tasks and datasets that rely less on identity recognition? To find out, in Experiment 3, we measured representational dissimilarities in humans (n=668) and CNNs on a completely different dataset using 60 images of distinct, non-famous young (approximate age between 20 and 30 years) male identities and a similarity-matching task (cf. Fig. 1B for the same task but on identity matching instead of similarity matching). We found the same pattern of results (Exp. 3; Fig. 2C; see Supplementary Note 4 for behavioral and CNN RDMs). Specifically, the face-identity trained CNN was again more similar to human behavioral similarities (Fig. 2C; Spearman’s *r*: 0.40) than the other three networks (all p=0, bootstrap test; Obj-Cat CNN *r*: 0.09, Obj-Face-Cat CNN r: 0.17, untrained CNN *r*: 0.05, all correlated with behavior above chance, all p<=.02). Taken together, face-identity trained networks, but not networks that were untrained or did not have training on face identification, represented faces similarly to humans (consistent with a recent study (30) testing view-invariant face representations), suggesting that optimization for face identification was necessary to match the human face representations tapped in behavioral judgements.

### Do CNNs show classic signatures of human face processing?

So far, we have found that CNNs optimized for face recognition achieve human-level face recognition performance and represent faces in a similar way, but CNNs optimized for object classification (even when trained extensively on face detection) do not. These findings suggest that human face recognition performance is unlikely to reflect a system optimized (through evolution, individual experience or both) generically for object categorization and/or face detection alone. Optimization for face recognition in particular seems to be required to capture human face recognition performance. But what about the classic behavioral signatures of human face processing, like the other-race effect and the face inversion effect? Why might human face recognition exhibit these phenomena? Might they also result from optimization for face recognition in particular? If so, we should expect to find that a CNN trained on face recognition, but not a CNN trained on object recognition (even if faces are included as an object category), would exhibit these same phenomena. In both cases, we predict the *presence* of the two signatures in the face-trained network, but not its magnitude, because the networks do not exactly match human experience.

In the other-race effect (34), humans show lower performance recognizing subgroups of faces they have had less exposure to during development. Previous work has suggested that CNNs also show such experience-dependent deficits when specific demographics are under-represented during training (35–37). But are these effects comparable to the other-race effect in humans, and could they result directly from optimization for recognition of faces of predominantly one race? Or is training of face detection or even passive exposure to faces of one predominant race (in Imagenet images during object classification training) sufficient? To find out, in Experiment 4, we tested white and Asian participants on a set of unfamiliar white and Asian female identities and compared them to CNNs using the same target-matching task (Fig. 1B). To test the other-race effect in CNNs, we trained a CNN on face recognition on a dataset of only Asian identities (Face-ID-Asian CNN), and another CNN on a predominantly white dataset with all Asian identities removed (Face-ID-white CNN). Further, we trained two networks on object categorization and face detection using only Asian identities (Obj-Face-Cat-Asian CNN) or only white identities (Obj-Face-Cat-white CNN).

Face recognition performance of the white participants (n=269) was lower for this Asian female test set (82.6%) set than for the white female test set (86.3%), replicating the other-race effect (Fig. 3A; light-gray bars; p=0, bootstrap test across participants). Importantly, Asian participants (n=102) showed the opposite, performing significantly better on the Asian test set (90.1%) than on the white test set (87.4%; p=0, bootstrap test). We found a significant interaction between the stimulus set and participants’ race (p=0, bootstrap test across participants), indicating that this effect cannot be simply due to differences in difficulty between stimuli sets. Our key question was whether this effect can be explained as a direct, perhaps inevitable, result of optimization for face recognition based on the demographically biased samples typical of human experience. Indeed, the recognition performance of the CNN trained on faces with Asian identities removed from the training set (Fig. 3A; Face-ID-white CNN) was significantly lower on Asian faces (88.8%) than on white faces (96.8%; p=0, bootstrap test), while the performance of the CNN trained on Asian faces (Face-ID-Asian CNN) was lower for white (86.6%) than for Asian face stimuli (91.5%; p=0, bootstrap test). Despite being matched in face experience, we did not find a significant difference in performance for the CNNs trained on object categorization and face detection of white identities (Obj-Face-Cat-white CNN: 63.1% white vs. 61.5% Asian; p=0.51, bootstrap test) or Asian identities (Obj-Face-Cat-Asian CNN: 64.8% white vs. 63.7% Asian; p=0.63, bootstrap test). Moreover, neither the CNN trained on object categorization nor the untrained CNN showed a significant drop in performance for the Asian set compared to the dataset of white female identities (Obj-Cat CNN: 62.5% white vs. 62.6% Asian; p=0.93, bootstrap test; Untrained CNN: 57.2% white versus 60.2% Asian; p=0.19, bootstrap test). Overall, these results suggest a reason why humans show an other-race effect: it is a natural consequence of training to discriminate faces from a specific race.

**Figure 3.**
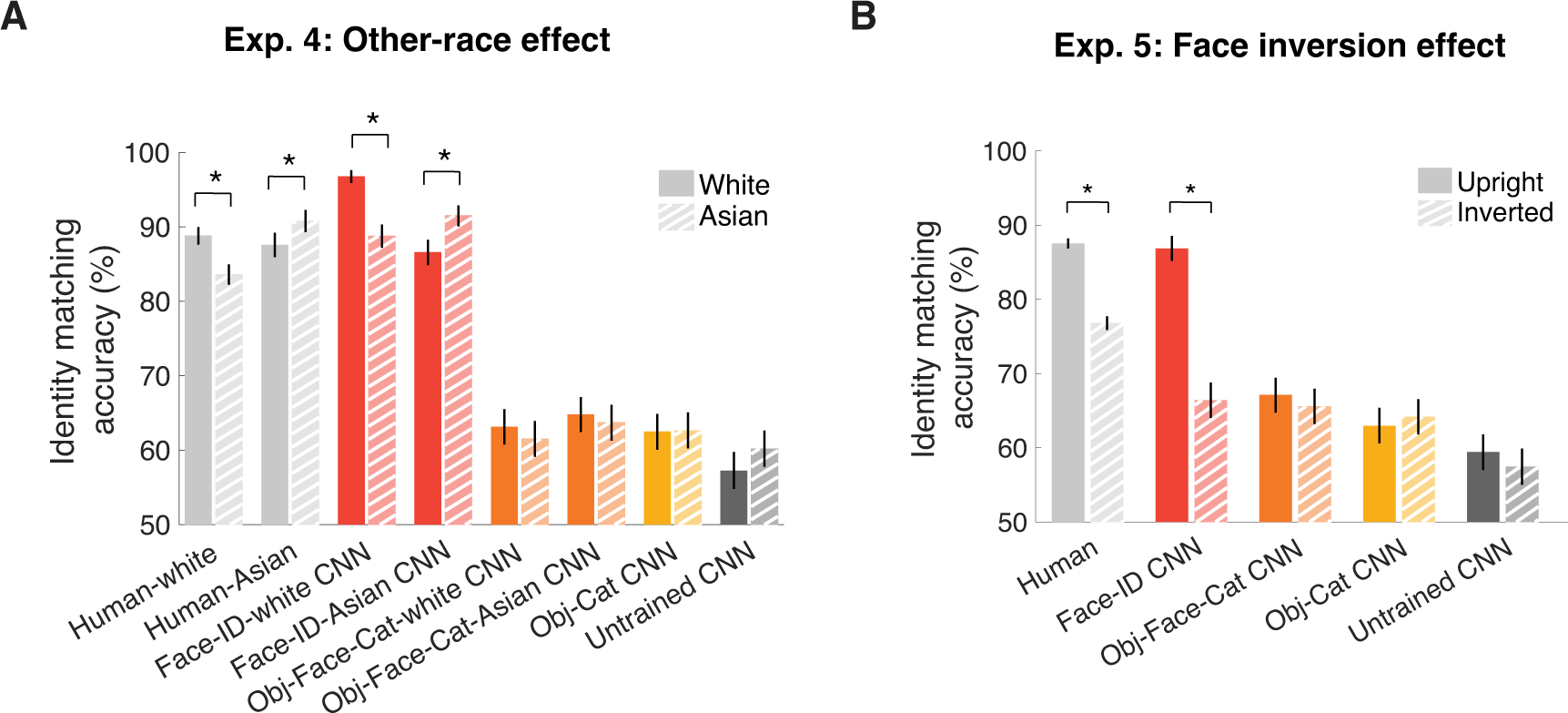
Face-identity trained CNNs show humanlike other-race effects (Experiment 4) and face inversion effects (Experiment 5). Face recognition performance was measured in a target-matching task (see Fig. 1B) in human participants (light gray), multiple face-trained CNNs (red), an object-and-face-trained CNN (Obj-Face-Cat CNN, orange), an object-trained CNN (Obj-Cat CNN, yellow) and an untrained CNN (Untrained CNN, dark gray). In CNNs, activations were extracted from the penultimate fully-connected layer to different stimuli sets and compared to human behavior (see Fig. 1B). **(A)** To measure the other-race effect, performance on unfamiliar, young female white identities (darker bars) was compared to performance on unfamiliar, young Asian female identities (lighter, striped bars). Human white participants (n=269) and the face-identity trained CNN (Face-ID-white CNN; with Asian identities removed from the training set), but not networks trained on object categorization and face detection of only white (Obj-Face-Cat-white CNN) or only Asian faces (Obj-Face-Cat-Asian CNN) or the object-trained or untrained CNN, showed significantly lower performance on Asian than white faces. In contrast, human Asian participants (n=102) and the CNN trained on Asian identities (Face-ID-Asian CNN) showed significantly lower performance on white than on Asian faces. Asterisks above bars indicate significant differences between conditions (*p=0, bootstrap test). **(B)** Identity matching accuracy for upright (n=1,532; darker bars) and inverted (n=1,219; lighter, striped bars) on white female identities from human behavior and CNNs. Only the face-identity trained CNN (Face-ID CNN) showed the face inversion effect, i.e. lower performance for inverted than upright faces, mirroring human behavior. Error bars denote bootstrapped 95% CIs. Asterisks above bars indicate significant differences between conditions (*p=0, bootstrap test).

Might optimization for recognition of (upright) faces similarly explain why humans show a face inversion effect? To test this prediction, in Experiment 5 we used the target-matching task with the white female identities used before (see Fig. 1B) but we presented them upside-down to both humans (n=1,219) and networks (Fig. 3B; see Fig. 4 for similar results using SVM decoding). Replicating multiple prior studies (3), human participants showed lower performance for inverted (76.8%) than for upright faces (87.5%; p=0; bootstrap test). This significant drop was also found as a significant within-participant difference using the subset of participants (n=364) who performed both the upright and inverted tasks (accuracy inverted: 75.9% versus upright: 87.5%; p=0, bootstrap test across participants). We found that the face-identity trained CNN (Fig. 3B; red) was the only network whose performance was lower for inverted than upright images (86.9% upright vs. 66.4% inverted; p=0, bootstrap test). Neither the object-categorization trained (Fig. 3B; yellow) nor the object-and-face-categorization trained (Fig. 3B; orange) or the untrained CNN (Fig. 3B; dark gray) showed a significant difference in performance between upright and inverted faces (Obj-Cat CNN: 63% upright vs. 64.2% inverted, p=0.30; Obj-Face-Cat CNN: 67.1% upright vs. 65.6% inverted, p=0.24 Untrained CNN: 59.4% upright versus 57.4% inverted; p=0.08, bootstrap test). Moreover, the face inversion effect was significantly larger for the face-identity trained than for all other CNNs (all p=0, bootstrap test). Thus, even though the object-trained network and the object-and-face-categorization trained network were exposed to faces, and much more to upright than inverted faces, neither shows the robust face inversion effect seen in the face-identity trained network. These findings show that the face inversion effect does not automatically arise from even extensive exposure or even training to detect upright faces, but it does result from optimization on (upright) face identification, providing a likely explanation of why humans show this effect.

**Figure 4.**
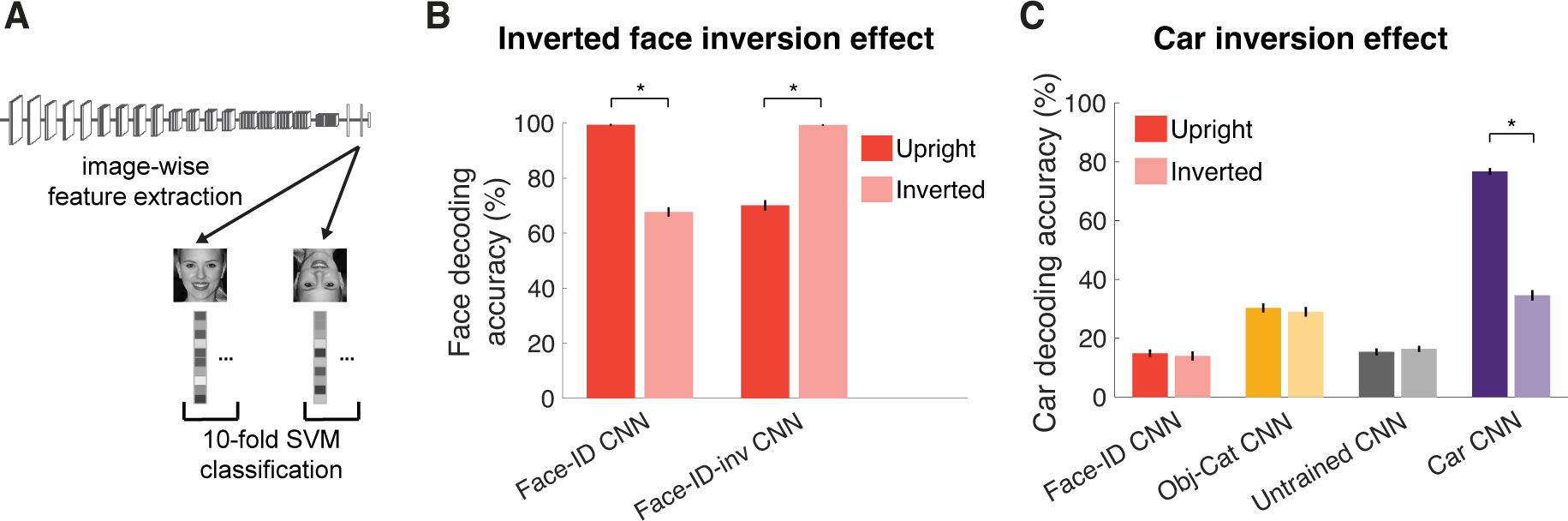
Inversion effects are not specific to upright faces per se. **(A)** Fine-grained category decoding accuracy was measured in CNNs trained on multiple tasks. Since there was no human data to match the performance to, we used a classification approach on larger stimulus sets to evaluate the performance in each condition. Activations were extracted from the penultimate fully-connected layer to different stimuli sets shown upright (darker bars) and inverted (lighter bars) and used to train and test a support vector machine (SVM). **(B)** To measure an “inverted face inversion” effect, 100 face identities (10 images each) were decoded from two face-identity trained CNNs: one trained on upright faces (Face-ID CNN, left), and one trained on inverted faces (Face-ID-inv CNN, right). While the CNN trained on upright faces showed higher accuracy for upright than inverted faces, the CNN trained on inverted faces showed the opposite. **(C)** We decoded 100 car model/make categories (10 images each) from the face-identity trained CNN (Face-ID CNN, red), the CNN trained on object categorization (Obj-Cat CNN, yellow), the untrained CNN (Untrained CNN, gray) and a CNN trained to categorize car models/makes (Car CNN, purple). Only the car-trained CNN showed an inversion effect for cars, i.e. lower performance for inverted than upright cars. Error bars denote SEMs across classification folds. Asterisks above bars indicate significant differences across conditions (*p<1e-5, two-sided paired t-test).

If indeed optimization for the recognition of upright faces is sufficient to produce a face inversion effect, hence explaining why humans show this phenomenon, does this reflect something special about face stimuli per se? Or, might any stimulus category produce inversion effects given sufficient training at fine-grained discrimination of exemplars within a category (2)? This question has long been pursued in the psychology literature (38), but the evidence that face-like inversion effects can result from perceptual expertise is mixed (39, 40). On the other hand, few if any humans have as much perceptual expertise on another stimulus category as they do for faces, and it remains unclear whether face-sized behavioral inversion effects might arise for non-face stimuli if they did. But with machine learning methods, we can give a network as much training on nonfaces as on faces. Here, we trained CNNs on inverted faces and on another fine-grained discrimination task (i.e., cars) to measure inversion effects in these networks and compared them to the previously used face-trained, object-trained and untrained CNNs. To evaluate the performance of these networks, we used a classification approach by training and testing a linear support vector machine (SVM) on activations for upright and inverted stimulus sets extracted from the penultimate fully-connected layer of these CNNs (Fig. 4A). The stimulus sets used here are larger (1,000 images) than those we used before because we were no longer constrained by the limitations of human experiments.

Indeed, we can produce an “inverted inversion effect” for faces by training the network only on inverted faces (Fig. 4B; Face-ID-inv CNN). That is, face decoding accuracy of 100 face identities (10 images each) from activations in the penultimate fully-connected layer of a CNN trained on inverted faces was significantly larger for inverted faces (99.3%) than for upright faces (70.1%; p<1e-5; two-sided paired t-test). In contrast, the decoding accuracy of the Face-ID CNN trained on upright faces showed the regular face inversion effect (Face-ID CNN: 99.4% upright vs. 67.7% inverted; p<1e-5). Note that the performance on the trained conditions (e.g., performance of upright faces for the Face-ID CNN) was larger than what we found before (i.e., in Fig. 1C). This difference could be due to differences in the stimulus sets or the different analysis method (i.e., SVM decoding approach) used here (see Table S3 for an overview), which allows for reweighting the features. Furthermore, we even find that a network trained on car discrimination shows a car inversion effect (Fig. 4C; Car CNN: 76.7% upright vs. 34.6% inverted; p<1e-5), but neither the object-trained (without cars included in the training set; Obj-Cat CNN: 30.3% upright vs. 29% inverted; p=0.66) nor the face-identity trained model (Face-ID CNN: 14.9% upright vs. 14% inverted; p=0.66) or the untrained model (Untrained CNN: 15.4% upright vs. 16.4% inverted; p=0.66) does. These findings indicate that inversion effects are not in principle specific to faces per se, but can in principle arise naturally for other stimulus classes from training on only upright stimuli.

## Discussion

Why does human face recognition show the particular behavioral signatures it does? Here we show that the characteristic human high accuracy, face space, other-race effects, and face inversion effects, are all found in CNNs optimized for face identification (on upright demographically biased training stimuli characteristic of human experience), but none of these effects are found in networks optimized for generic object categorization. Further, none of these effects arise in networks that are trained on face detection only, despite having the same amount of face experience as the face-identity trained networks. This finding shows that face experience alone is not sufficient to produce these effects. Rather, it is optimization for the specific task of discriminating individual faces from each other that produces these effects. These findings enable us to go beyond the mere documentation of the “special” signatures of the human face system, to provide an answer to the question of *why* human face recognition exhibits these phenomena. What our findings suggest is that these classic behavioral signatures of face recognition in humans result from optimization for face recognition in particular. Thus we might expect any system optimized on this task to show the same phenomena.

We further find that the most classic signature of the face system, the face inversion effect, need not in principle be restricted to face stimuli. In CNNs trained on cars, we find a car inversion effect, and in CNNs trained only on inverted faces, we find an inverted inversion effect. This kind of behavioral inversion effect for nonface objects of expertise has long been sought experimentally in humans, but the enterprise has remained inconclusive (38–40), probably because it is very difficult to find any stimulus class for which humans have as much expertise as they do for faces. With CNNs however, we can control exactly how much experience and what task each network is trained on. These methods have enabled us to show that we should not expect faces to be “special” in the kinds of representations we extract from them. Rather, faces are special in the human brain (41), and in networks jointly trained of face and object recognition (42) in that distinct neurons and network units are engaged in their processing.

Our results also give some hints about the possible origins of the other-race effect. Like our human participants, we found that the CNN trained on predominantly white faces (Asian identities were removed from the training set) showed a drop in performance for Asian faces. This finding mirrors recent reports of bias in AI face recognition systems (35–37, 43) and suggests a computational account of the other-race effect in humans (34). Thus, achieving high face recognition accuracy in machines (and possibly also humans) requires not only extensive face experience, but extensive experience within each of multiple face types. This finding, along with our finding that the face inversion effect arises spontaneously in CNNs trained to discriminate face identities but not in CNN trained on face detection and/or object classification, accords with other findings showing signatures of human face perception in face-identity trained networks, such as face familiarity effects (22), the Thatcher illusion (44) and view-invariant identity representations (30, 45, 46).

Of course, CNNs differ from brains in myriad ways, perhaps most strikingly in how they learn. We are not arguing that the human face system develops in the same way CNNs do, certainly not from extensive labelled examples trained with backpropagation (47). Rather, our point is that CNNs allow us to move beyond the mere documentation of behavioral signatures of face processing as curiosities to be collected, to the more interesting enterprise of asking which of these signatures may be explained as a consequence of the computational optimization for face recognition (6, 48).

Despite the consistency of our results with CNNs trained on face recognition, many puzzles remain. For example, given our finding that face-identity trained networks better explain human face perception, why do object-trained CNNs do similarly or even better at explaining face-specific neural responses (16, 49)? One possible explanation is that human face behavior might be read out from later stages of neural processing than have been investigated so far in studies examining the correspondence between CNNs and neural responses. This hypothesis is supported by several findings suggesting that face-specific regions in the superior temporal sulcus (50, 51), or areas beyond the core system of face perception (52, 53) may be involved in face identity recognition in humans or monkeys (54). Another explanation could be that the resolution of neuroimaging methods in humans is insufficient to read out identity information in face-specific areas (55). However, methodological limitations of fMRI cannot fully explain this discrepancy, because face-trained models also did not outperform object-trained models in predicting human intracranial data (16), which provides higher spatial resolution than fMRI. A third explanation could be that object-specific features can be repurposed for face perception by reweighting the features (as is typically done when building encoding models). However, we recently found that even when training a support vector machine (SVM) on object-specific features, those features were much less useful for face identification than face-trained features (42). Lastly, neural face representations might be optimized for fine-grained face discrimination and face detection. While standard face-trained CNNs are trained on to discriminate different faces from each other, the face-specific features they develop were not optimized to distinguish faces from objects. This hypothesis could be addressed by training a CNN on fine-grained face recognition and object categorization simultaneously. Indeed, our recent work suggests that networks optimized for both face and object recognition spontaneously segregate face from object processing in the network, and are able to capture human behavioral representational space for faces and objects (42). It will be of interest to directly compare this network to neural responses. In the future, these questions might be answered by combining human behavior, neural data and deep neural networks to find out which task optimization best explains neural face responses and where in the brain the face representations tapped in behavioral tasks reside.

Would more complex types of networks better match human face perception? While it is possible that recurrent neural networks (56, 57) or 3D-based generative models of face perception (49, 58–60) could also explain our data, it is unlikely that they will outperform face-identity trained CNNs given the high consistency we observed (sometimes nearly reaching noise ceiling) between CNNs and human face perception behavior. This finding suggests that simple feedforward CNNs are sufficient to model behavioral face signatures. However, feedforward CNNs do not perform as well on all face tasks (e.g., the Hollow-face effect (58)) and it will be critical to study these tasks further. Moreover, our task was designed to test face recognition under relatively high image variation conditions, but it remains possible that tests with even higher image variation would reveal a gap between humans and feed-forward networks.

Our work also provides some clues into the origins of the human face system, by showing that humanlike face recognition can in principle arise from face-specific experience alone, but only if networks are trained to discriminate individual faces from each other. Importantly, however, training on object classification alone, even with extensive experience on face detection appears not to be sufficient. This finding highlights the fact that the behavior of a network depends not only on the training diet, but also on the training task. It remains an open question whether the relevant face experience that shaped the human face recognition system occurred during evolution, or modern individual experience, or (as is usually the case) both.

In sum, our findings, that face-identity trained but not object-trained models, even when trained on face detection with the same amount of face experience, match many of the classic signatures of human face recognition enable us to explain these signatures of human face processing as the expected result of optimization for this specific task. Our study joins several other recent investigations that use deep neural networks to explain human perceptual phenomena as the result of optimization for a given task (6, 48). For example, many phenomena in auditory processing were mirrored in networks trained on natural sounds (61, 62). Moreover, the existence of special neural populations selectively engaged in face recognition can also be explained as the result of joint optimization for face and object recognition (42). Each of these studies uses CNNs to move beyond the mere characterization of perceptual phenomena to address the more fundamental question of why our perceptual systems work the way they do. This new strategy builds upon earlier work using ideal observer methods in perception, but enables us to now tackle more complex real-world perceptual problems.

## Materials and Methods

### Experiment 1: Comparing human face recognition performance in the target-matching task to task-optimized CNNs

#### Participants

A total of 1,540 individual workers from the online crowdsourcing platform Amazon Mechanical Turk participated in the target-matching tasks (Figure 1b) on white upright stimuli. A total of 8 workers were excluded from the analysis due to overly fast responses (response time in more than five trials < 500ms or more than 10 trials <800 ms). All workers were located in the United States. We asked workers to voluntarily provide their sex, race and age (i.e., in ranges ‘18-24’, ‘25-34’, ‘35-44’, ‘45-54’, ‘55-64’, ‘65 or older’). The average workers’ age was between 25 and 34 years, 57% of workers were female, 42% were male and 1% reported ‘other’ or did not report their sex. The majority of the workers were white (70%), 15% were Black, 10% were Asian and 5% reported ‘other’ or did not report their race. To avoid familiarity effects with identities, each worker was only allowed to perform one set of 21 trials (using all 40 distinct identities) per task. Some workers were still able to perform more sets of trials due to technical restrictions on Amazon Mechanical Turk. In this case, only the first set of trials was included in the analysis. The number of workers was chosen such that each trial was sampled 20 times across workers. This number was chosen based on previous studies that sampled triplets on Amazon Mechanical Turk (63).

#### Stimuli and behavioral target-matching task on upright faces

To measure human behavioral face recognition performance, participants performed a target-matching task on Amazon Mechanical Turk. To construct this task, we chose 5 images of each of 40 identities. To ensure that the task would not merely rely on external face dimensions (e.g., age, gender), we restricted the stimuli to white female identities of similar age (approximately 20-35 years). We further tried to choose individuals who were less famous to reduce familiarity effects. Moreover, we chose images of these identities that varied across lighting, hairstyle, pose and other low- and mid-level image features (as validated by the low performance of early convolutional layers on this task, around 60% correct; see Supplementary Note 1). We first selected identities which fulfilled these criteria from the Labeled Faces in the Wild dataset (64). Since the Faces in the Wild dataset did not contain sufficient identities to fulfill these criteria, we manually supplemented more identities by selecting them from the internet and by using identities from the VGGFace2 dataset. We then randomly chose images to build triplets in which each target identity (2 images) was paired with each other identity as distractor (1 image) for a total of 1560 (40×39) triplets. Critically, none of the test identities were used for training the Face CNN. Also importantly, from the 156,000 possible triplets (40 identities x 5 images x 4 images (same identity) x 195 images (distractor identity)) available, we only used the 1560 triplets that were presented to the human participants to compute the identity recognition performance of CNNs. On Amazon Mechanical Turk, we asked human participants to choose which of two face images (e.g., matches A or B) belonged to the same identity as the third target image. The position of the matching images (left or right) was pseudo-randomized across trials. To reduce perceptual and behavioral noise as much as possible, participants had unlimited time to perform each trial. Each participant performed 20 trials in which 20 distinct identities were paired with the remaining half of the 20 identities such that each identity was shown only once during the set of trials. To measure within-participant reliability, participants also performed an additional trial in which one randomly chosen trial from the initial set of 20 trials was repeated. Each triplet was repeated 20 times across participants and the average identity matching performance across all triplets and human participants served as measure for human face recognition performance.

#### Untrained and trained convolutional neural networks

To explore the role of face experience in face recognition, we trained four CNNs with varying amounts and kinds of visual experience. All CNNs were based on the VGG16 architecture (24) (see Supplementary Note 2 for parallel results based on Alexnet (27) and ResNet (28, 29) architectures). First, to test the performance of an untrained network, we used a randomly-initialized VGG16 network (Untrained CNN). Second, to measure the performance on face recognition on a CNN trained on generic objects, we trained a randomly-initialized VGG16 architecture on object categorization only (Obj-Cat CNN). The Object CNN was trained on 423 manually sampled object categories of the ILSVRC-2012 database (65). To obtain an object-trained CNN, we removed all categories from the original dataset that were not prototypical objects (e.g., scene-like categories, animals). The remaining 423 categories consisted of prototypical objects (e.g., trumpet, hammer, coffee cup). For each of the 423 selected object categories, we used 1000 images for training and 50 for validation, for a total of 423k training and ∼21k validation images. Third, we trained a VGG16 network on face identity categorization only (Face-ID CNN). The network was trained on 1,714 identities (857 female) from the VGGFace2 database (8). Specifically, we chose identities with a minimum of 300 images per identities but otherwise the identities were randomly selected from the VGGFace2 database. Importantly, to allow for better comparison with humans, we further tried to match the number of identities to the minimum number of identities typically known by humans (range 1,000 – 10,000) (66). To best match the training set size to the object-trained CNN, we randomly chose 246 images per face identity for training, and 50 images per category for validation, for a total of ∼422k training and ∼85k validation images. Fourth, to test whether mere face experience (without discriminating individual faces) was sufficient to match human face processing, we included a CNN that was matched in the amount of face experience to the face-identity trained CNN, but trained on coarse face detection only. Specifically, we trained a CNN on object and face categorization (Obj-Face-Cat CNN) on the exact same 423 object categories as the object-trained CNN with one additional category (i.e., 424 categories in total) that included all face images used to trained the face-identity trained CNN. Note, that this network did not only include the same amount of face images as the face-identity trained model, but was also trained on a much larger training set.

#### Target-matching task on upright faces in CNNs

To directly compare the face recognition performance between humans and CNNs, for the behavioral target-matching tasks on face images, we presented the same stimuli to the four different CNNs: the untrained CNN, the object-categorization-trained CNN, the object-and-face-categorization CNN and the face-identification CNN. For each tasks and stimuli, we simulated the behavioral task in CNNs by extracting activation from the penultimate fully-connected layer to each of the 200 images (see Supplementary Note 1 for additional layers). We measured the pairwise similarity for each pair of images using correlation distance (1 – Pearson’s *r*) between the activation patterns. The network’s choice was modeled as which of the two matching images was closest to the target image. Critically, from the 156,000 possible triplets (40 identities x 5 images x 4 images (same identity) x 195 images (distractor identity)) available, we only used the 1,560 triplets that were presented to the human participants to compute the identity recognition performance. Note that when we compared CNNs to participants who indicated that they were unfamiliar with all identities, we only compared the performance on the triplets performed by those participants. By averaging the choice accuracy (0 or 1 for incorrect or correct, respectively) across all 1,560 triplets, we obtained a corresponding identity matching performance for each of the different CNNs.

### Experiment 2: Using a multi-arrangement task to compare human perceptual similarity of faces to task-optimized CNNs

#### Participants

Behavioral data from 14 laboratory participants (7 female; mean age 25.9, SD = 4.33) from a previously published study (67) were used to perform the representational similarity analysis (RSA) using the multi-arrangement task. As described previously, all participants provided informed, written consent prior to the experiment and were compensated financially for their time. The Massachusetts Institute of Technology (MIT) Committee on the Use of Humans as Experimental Subjects approved the experimental protocol (COUHES No 1606622600).

#### Stimuli and behavioral representational dissimilarities

To find out whether humans and CNNs represent faces similarly, we performed RSA in two different experiments. The experimental design and stimuli to obtain the behavioral data have been explained in detail previously (67), so here we just briefly summarize the stimuli and task. Participants performed a multi-arrangement task (33) using 80 face stimuli. Stimuli consisted of 5 images of each of 16 celebrities, which varied orthogonally in gender and age, such that half were female and half were male and half of them were young (below ∼35 years) and half were old (above ∼60 years). Participants performed the multi-arrangement experiment online using their own computer. During the task, participants were instructed to arrange different subsets of the images based on their perceived similarity (“similar images together, dissimilar images apart”) by dragging and dropping them in a circle. After the completion of the experiment, the pairwise squared on-screen distances between the arranged images was computed, thus representing a behavioral representational dissimilarity matrix (RDM; see Fig. 2A bottom left for visualization of the mean behavioral RDM). For each participant, we extracted the lower off-diagonal data from the behavioral RDM to obtain a vector of pairwise dissimilarities used for computing the correlations.

We additionally computed the noise ceiling for the representational dissimilarities given the inconsistencies across participants using a method described previously (68). Briefly, we estimated the upper bound of the noise ceiling as the mean correlation of each participant’s vector of perceived dissimilarities with the group mean (including the participant itself). In contrast, the lower bound was computed by taking the mean correlation of each participant with all other participants.

#### Representational similarity analysis between humans and CNNs

To obtain representational dissimilarities in CNNs, we presented the same stimuli as used for the human participants to the four CNNs. For each CNN, we extracted the activation patterns to each image separately from the penultimate layer (see Supplementary Note 5 for other layers) and computed the correlation distance (1 – Pearson’s *r*) between each pair of activation patterns. This resulted in one RDM for each of the four CNNs (see Supplementary Note 4 for visualization of the RDMs).

To compute the similarity between the human RDMs and the RDMs obtained for the CNNs, we rank correlated each participant’s behavioral dissimilarities vector with the corresponding CNN dissimilarities vectors. The average rank correlation across participants served as similarity measure between human participants and CNNs. We further computed the bootstrapped confidence intervals by bootstrapping the participants and computing the correlation with the CNN RDMs 10,000 times and computed the 95% CI of the resulting distribution.

#### Statistical inference

To measure statistical significance, we used bootstrap tests. Specifically, we bootstrapped the participant-specific RDMs 10,000 times and correlated them with the CNN RDMs obtained from the penultimate fully-connected layer of each network to obtain an empirical distribution of the correlations. The standard deviation of these distributions defined the SEM for the correlation between humans and CNNs. To test for differences (or differences of differences) between correlations, we bootstrapped the participants 10,000 times and computed the mean difference between correlations resulting in an empirical distribution of correlation differences.

### Experiment 3: Using a similarity-matching task to compare human perceptual similarity of faces to task-optimized CNNs

#### Participants

A set of 697 individual workers participated in the similarity-matching task (Fig. 2C) on Amazon Mechanical Turk. A total of 29 workers were excluded from the analysis due to overly fast responses (response time in more than five trials < 500ms or more than 10 trials < 800 ms). All workers were located in the United States. The average workers’ age was between 25 and 34 years, 56% of workers were female, 42% were male and 2% reported ‘other’ or did not report their sex. The majority of the workers were white (71%), 19% were Black, 8% were Asian and 2% reported ‘other’ or did not report their race. For this task, workers were not restricted in the number of trials they could perform. All workers provided informed consent and were compensated financially for their time. The experimental protocol was approved by the Massachusetts Institute of Technology (MIT) Committee on the Use of Humans as Experimental Subjects (COUHES No 1806424985) and conducted following all ethical regulations for conducting behavioral experiments.

#### Stimuli and behavioral representational dissimilarities

To test whether the results from the multi-arrangement task from Experiment 2 would generalize to a different dataset and task, we conducted a similarity-matching task on Amazon Mechanical Turk. To construct this task, we chose one image of each of 60 unfamiliar male identities from the Flickr-Faces-HQ database (69). The stimuli included 60 young male identities of similar age (approximately between 20 and 30 years old) with a neutral facial expression. Participants were asked to choose which of two images was more similar to a third target image (i.e., triplet). Each of the possible triplets (60×59×58/2 for a total of 102,660 triplets) was sampled once (although some triplets were excluded, see Participants). The choice for a specific triplet resulted in two pairwise similarities: between the target and each of the matching images (e.g., when the choice for the triplet with target A and matches B and C was C, this would result in “1” for the pair A-C, and “0” for the pair A-B). Thus, the perceived similarity of each pair of face images was on average sampled ∼100 times (i.e., each of the 102,660 triplets produced two of the 1,770 (60×59/2) pairwise similarities resulting in 116 (102,660/1,770) samples without exclusions). The proportional number of times that each target-match pair was chosen as the more similar pair was used as similarity value for the pair and converted into a dissimilarity value by subtracting it from 1. We extracted the lower triangle excluding the diagonal from the resulting dissimilarity matrix (see Supplementary Note 4 for visualization of the behavioral RDM) to obtain a vector of pairwise dissimilarities. We then used the vector of dissimilarity values to compute the similarity with the CNNs.

To compute the noise ceiling in this task, we used split-half correlation. Since participants were not limited in the number of trials they could perform, participants contributed with varying degrees to the final dissimilarity matrix. To not bias the split-half correlation by participants who contributed more trials, we only used the first set of trials collected by each individual participant. We randomly split the participants into two halves 50 times and computed the correlation between the dissimilarity vectors based on the two halves for each split. We then used the mean correlation added and subtracted by twice the standard deviation of this set of correlations as noise ceiling.

#### Representational similarity analysis between humans and CNNs

As in Experiment 2, we obtained representational dissimilarities in CNNs, by presenting the same stimuli used for the human participants to the four CNNs. For each CNN, we extracted the activation patterns to each image separately from the penultimate layer (see Supplementary Note 5 for other layers) and computed the correlation distance (1 – Pearson’s *r*) between each pair of activation patterns. This resulted in one RDM for each of the four CNNs (see Supplementary Note 4 for visualization of the RDMs).

To compute the similarity between the human RDMs and the RDMs obtained for the CNNs, we rank-correlated the human behavioral dissimilarity vector obtained from the pairwise proportional choices across all triplets with the corresponding CNN dissimilarity vector constructed from the same stimuli.

#### Statistical inference

To measure statistical significance, we used the same bootstrap tests as in Experiment 2, but bootstrapped the dissimilarity vectors instead of participants. Specifically, we bootstrapped the dissimilarity values of the behavioral RDM and CNN dissimilarity vectors 10,000 times and computed the rank correlation to obtain 95% CIs, and to compute a distribution of correlation differences. All p-values were derived as explained below (see section Statistical Inference).

### Experiment 4: Comparing the other-race effect between humans and task-optimized CNNs

#### Participants

To study the other-race effect (Fig. 3A), we collected data using a different image dataset. Specifically, we collected data on unfamiliar, young white and Asian female stimuli using the target-matching task. To sample white participants (n=396), we used Amazon Mechanical Turk and only included workers who were located in the United States, who listed their race as white and who reported that during elementary school at least 50% of their peers were white and less than 50% were Asian. Of those workers, 127 had to be excluded due to overly fast responses, overly deterministic responses (more than 65% left or right clicks only, when chance level was 50%) or because their response differed in the catch trial. The average workers’ age was between 25 and 44 years, 46% of workers were female, 53% were male and 1% reported ‘other’ or did not report their sex. Amazon Mechanical Turk does not provide access to workers that are based in East-Asian countries. Therefore, to additionally collect Asian participants on the same task, we used Clickworker (www.clickworker.com; which provides access to workers from some countries in East Asia) to recruit participants and directed them to perform the experiment on the Meadows platform (www.meadows-research.com). We were able to recruit 132 participants who listed their race as Asian and who reported that during elementary school at least 50% of their peers were Asian. Of those participants 30 had to be excluded using the same exclusion criteria as for the white participants. The average workers’ age was between 35 and 44 years, 44% of workers were female, 54% were male and 2% reported ‘other’ or did not report their sex.

All workers provided informed consent and were compensated financially for their time. The experimental protocol was approved by the Massachusetts Institute of Technology (MIT) Committee on the Use of Humans as Experimental Subjects (COUHES No 1806424985) and conducted following all ethical regulations for conducting behavioral experiments.

#### Stimuli and behavioral target-matching task on white and Asian faces

To test the other-race effect in humans and compare them to CNNs (see section below), we ran the same test on another set of face stimuli: unfamiliar, young, female white and Asian faces. We recruited white human participants (n=269) using Amazon Mechanical Turk, only including participants who were white and who reported that more than 50% of their peers were white during elementary school. Due to restriction of workers on Amazon Mechanical Turk, we had to recruit participants from Asia (n=102) using Clickworker. Once they signed up for the experiment on Clickworker, participants were forwarded to the Meadows platform running the same task as our white participants on Amazon Mechanical Turk. To exclude that familiarity with some of the identities would influence the results, we collected a novel set of 5 images of each of 80 identities (40 of each race) by using photos provided by colleagues, and by sampling photos of identities on Instagram (with less than 2,000 followers). All of the identities were female and between 20-35 years of age, and none of them was used as training for the CNNs. For each race, we built triplets in which each target identity (2 images) was paired with each other identity as distractor (1 image) for a total of 1560 (40×39) triplets. During the experiment, each participant performed 20 trials of each race, randomly interleaved, for a total of 40 trials. For each participant, 20 distinct identities of a race were paired with the remaining half of the 20 identities of the same race, such that each identity was shown only once during the set of trials. To measure within-participant reliability, we included an additional trial in which one randomly chosen trial from the set of 40 trials was repeated. All other experimental parameters were identical to the target-matching task described above.

#### Testing the other-race effect in CNNs

To test the other-race effect in CNNs, we trained four additional CNNs. We first trained a VGG16 architecture on a dataset of mainly white identities (Face-ID-white CNN). To obtain such a dataset, we manually removed all Asian identities from the identities we previously selected from the VGGFace2 dataset. Specifically, we removed 60 Asian identities from the set of 1,714 identities, for a total of 1,654 remaining mainly white identities. We then trained another VGG16 network on Asian identities (Face-ID-Asian CNN) using the Asian Face Dataset (AFD (70)). We randomly chose 1,654 identities of this dataset to match the number of identities of the Face-ID-white CNN. These identities had a minimum of 105 images per identity (∼174k images). To avoid imbalanced classes, we therefore chose 105 images for each of the identities of both datasets using 100 for training and five images for validating. To further test whether training on face detection would be sufficient for the other-race effect to emerge in CNNs, we additionally trained two VGG16 networks on object categorization and face detection using the white identities only (Obj-Face-Cat-white) or the Asian identities only (Obj-Face-Cat-Asian).

We then presented the same unfamiliar white and Asian face stimuli as used during the behavioral tasks to the four trained CNNs as well as the Obj-Cat CNN and the untrained CNN and extracted activations from the penultimate fully-connected layer (see Supplementary Note 1 for other layers). All other analyses were identical to the analyses to the target-matching task on upright face images as described above. To test whether the other-race effect was restricted to a particular architecture, we further investigated the other-race effect in Alexnet and Resnet architectures trained on face identification or object categorization or untrained (see Supplementary Note 2).

### Experiment 5: Comparing the face inversion effect in humans and task-optimized CNNs

#### Participants

In addition to the set of workers participating the target-matching task on upright stimuli in Experiment 1, a total of 1,237 individual workers from the online crowdsourcing platform Amazon Mechanical Turk participated in the target-matching tasks (Fig. 1B) on inverted stimuli. A total of 18 workers were excluded from the analysis due to overly fast responses (response time in more than five trials < 500ms or more than 10 trials <800 ms). Each worker was only allowed to perform one set of 21 trials (using all 40 distinct identities) per task. Some workers were still able to perform more sets of trials due to technical restrictions on Amazon Mechanical Turk. In this case, only the first set of trials was included in the analysis. Of the remaining workers, 64 workers had also participated in the target-matching task of upright face images (Experiment 1). In addition to these 64 workers, 300 of the workers on Amazon Mechanical Turk that participated in the target-matching task on inverted images (independent of the 64 workers who had also participated in Experiment 1) performed the same task on upright images. In total, we recruited 364 workers on the target-matching task using both upright and inverted images.

All workers provided informed consent and were compensated financially for their time. The experimental protocol was approved by the Massachusetts Institute of Technology (MIT) Committee on the Use of Humans as Experimental Subjects (COUHES No 1806424985) and conducted following all ethical regulations for conducting behavioral experiments.

#### Stimuli and behavioral target-matching task on inverted stimuli

To measure human behavioral face recognition performance on inverted faces, participants performed the same target-matching task on Amazon Mechanical Turk as described above. We used the identical stimuli as for the target-matching task on upright faces, but presented them upside down.

#### Target-matching task using inverted face stimuli in CNNs

To directly compare the face recognition performance between humans and CNNs, for the behavioral target-matching tasks on upright and inverted white face images, we presented the inverted stimuli to the four different CNNs (Face-ID CNN, Obj-Face-Cat CNN, Obj-Cat CNN, Untrained CNN) and extracted activations from the penultimate fully-connected layer (see Supplementary Note 1 for other layers). All other analyses were identical to the analyses described for the target-matching task on upright face images above.

To test whether the face inversion effect was restricted to a particular architecture, we further investigated the face inversion effect in Alexnet and Resnet architectures trained on upright face identification or object categorization or untrained (see Supplementary Note 2).

### Statistical inference

For all analyses, we used non-parametric statistical tests that do not rely on assumptions about the distributions of the data. For the target-matching tasks, we bootstrapped the combination of paired identities shown as a trial (i.e., 1,560 triplets) 10,000 times and averaged the responses to obtain a distribution of accuracies. The 2.5^th^ and the 97.5^th^ percentile of this distribution were used as 95% confidence interval (CI) for the behavioral and CNN performances. For statistical inference of the differences between performances, we bootstrapped the triplets 10,000 times and computed the mean difference between accuracies resulting in an empirical distribution of performance differences. To test for differences between differences (i.e., interaction effects), we performed the same analysis but bootstrapped the difference of the differences 10,000 times. The number of differences (or differences of differences) that were smaller or larger than zero divided by the number of bootstraps defined the p-value (i.e., two-sided testing). In the case of within-participant tests, we performed the same analysis but bootstrapped the participants (instead of triplets) 1,000 times to obtain a distribution of performance differences or difference of differences in case of interaction effects. All p-values were corrected for multiple comparisons using false discovery rate (FDR) at a 0.05 level.

### Extended testing for inversion effects in task-optimized CNNs

#### Training convolutional neural networks

To test whether a CNN trained only on inverted faces (Face-ID-inv CNN) would show an inverted face inversion effect, we additionally trained a CNN on inverted face images. We used the same architecture, training dataset and parameters as for the Face-ID CNN but showed all images inverted instead of upright during training.

Further, to test whether inversion effects could emerge in CNNs trained on a different domain of stimuli, we trained a VGG16 architecture on car model/make discrimination (Car CNN) using the CompCars dataset (71). To obtain enough images per class, we concatenated images of this dataset from the same model/make but of different years into one class. In this fashion, we ended up with 1109 classes with 45 images for training and 5 images for validating per class (∼50k training and ∼5.5k validation images).

#### Decoding of visual categories in CNNs

To test whether CNNs trained on varying tasks differ in how much information about a visual category they contain, we decoded exemplars of independent sets of visual categories from activations extracted from those networks (Fig. 4A).

To test the inverted face inversion effect (Fig. 4B), we used 100 held-out face identities (50 female; 10 images per identity; 1000 images in total) from the VGGFace2 dataset that were not included in the training set of the CNNs. We presented these images upright and inverted and extracted activations from both the CNN trained on upright faces (Face-ID CNN) and the CNN trained on inverted images (Face-ID-inv CNN).

To test whether the inversion effect was specific to faces, we further tested for an inversion effect for fine-grained car decoding (Fig. 4C). We selected 10 images of 100 model/make categories from the CompCars dataset (1000 images in total) that were not including in the training of any network and extracted activations to those images from the face-identification trained CNN, the object-categorization trained CNN (which had no vehicle-related categories in the training set), the untrained CNN and a CNN trained on car discrimination.

For both of these analyses, we extracted the activation in the penultimate layer of each network (i.e., the last layer before the classification layer) to the image sets. For each task and activations from each network, we trained and tested a 100-way linear support vector machine (with L2 regularization) on the corresponding activation patterns using a leave-one-image-out (i.e., 10-fold) cross-validation scheme. We computed the mean and SEM across classification folds and used two-sided paired t-tests across classification folds to test for differences between decoding accuracies.

### Training parameters for CNNs

For all trained networks (i.e., Face-ID, Face-ID-white, Face-ID-Asian, Face-ID-inv, Obj-Face-Cat, Obj-Face-Cat-white, Obj-Face-Cat-Asian, Obj-Cat, Car CNN), we used similar training parameters as suggested in (24): stochastic gradient descent (SGD) with momentum with an initial learning rate of 10^-3^, a weight decay of 10^-4^ and momentum of 0.9. We trained each network for at least 50 epochs (i.e., full passes over the training set) and the learning rate was reduced twice when the training loss saturated to 10^-4^ and 10^-5^, respectively. All CNNs were trained until the training loss reached saturation before being compared to human data. To update the weights during training, we computed the cross-entropy loss on random batches of 128 images and backpropagated the loss. Each image was scaled to a minimum side length (height or width) of 256 pixels, normalized to a mean and standard deviation of 0.5. For data augmentation, we showed images randomly as gray-scale with a probability of 20% and chose random crops of the size of 224 x 224 pixels (out of the 256 x 256 pixel-sized images) for each image during training. All training parameters were selected in pilot experiments. The test images were scaled, normalized and center-cropped before extracting the activation patterns.

### Data availability

All stimuli used for the online experiments are available at https://osf.io/dbks3/. The stimuli used for the laboratory experiment have been previously made available at https://osf.io/gk6f5/. The source data underlying Figures 1-4 and Supplementary Figures 1-10 will be made available on OSF upon acceptance.

### Code availability

To reproduce the relevant analyses and figures, the Python and Matlab scripts will be made available on OSF upon acceptance. The code to train computational models has been previously made available at https://github.com/martinezjulio/sdnn.

## Acknowledgements

We thank Martin Hebart for his help and advice in setting up the experiments on Amazon Mechanical Turk, Rebecca Saxe for valuable comments on the manuscript, Elizabeth Mieczkowski for help with online experiments and members of the Kanwisher lab for fruitful discussions and feedback. This work was supported a Feodor-Lynen postdoctoral fellowship of the Humboldt Foundation to K.D., the Deutsche Forschungsgemeinschaft (DFG, German Research Foundation; project number 222641018–SFB/TRR 135), “The Adaptive Mind”, funded by the Excellence Program of the Hessian Ministry of Higher Education, Science, Research and Art to K.D., NIH grant Grant DP1HD091947 to N.K and National Science Foundation Science and Technology Center for Brains, Minds, and Machines.

## References

1. M. J. Farah, Is face recognition ‘special’? Evidence from neuropsychology. Behav Brain Res 76, 181–189 (1996).

2. R. Diamond, S. Carey, Why Faces Are and Are Not Special: An Effect of Expertise. J. Exp. Psychol. Gen. 115, 107–117 (1986).

3. R. K. Yin, Looking at upside-down faces. J. Exp. Psychol. 81, 1–5 (1969).

4. R. K. Bothwell, J. C. Brigham, R. S. Malpass, Cross-racial identification. Personality and Social Psychology Bulletin 15, 19–25 (1989).

5. T. Valentine, “Face-space models of face recognition” in Computational, geometric, and process perspectives on facial cognition: Contexts and challenges., M. J. Wenger, J. T. Townsend, Eds. (2001), pp. 83–113.

6. A. J. Kell, J. H. McDermott, Deep neural network models of sensory systems: windows onto the role of task constraints. Curr. Opin. Neurobiol. 55, 121–132 (2019).

7. N. Kanwisher, M. Khosla, K. Dobs, Using Artificial Neural Networks to ask Why Questions of Minds and Brains. Trends Neurosci. (2022).

8. Q. Cao, L. Shen, W. Xie, O. M. Parkhi, A. Zisserman, VGGFace2: A dataset for recognising faces across pose and age in IEEE International Conference on Automatic Face & Gesture Recognition, IEEE International Conference on Automatic Face & Gesture Recognition., (2018), pp. 67–74.

9. Y. Taigman, M. Yang, M. A. Ranzato, L. Wolf, DeepFace: Closing the Gap to Human-Level Performance in Face Verification in Proceedings of the IEEE Conference on Computer Vision and Pattern Recognition (CVPR), Proc. Computer Vision and Pattern Recognition., (2014), pp. 1701–1708.

10. P. J. Phillips, et al., Face recognition accuracy of forensic examiners, superrecognizers, and face recognition algorithms. Proc. Natl. Acad. Sci. U.S.A. 115, 6171–6176 (2018).

11. R. Zhang, P. Isola, A. A. Efros, E. Shechtman, O. Wang, The Unreasonable Effectiveness of Deep Features as a Perceptual Metric in Proceedings of the IEEE Conference on Computer Vision and Pattern Recognition (CVPR), Proceedings of the IEEE Conference on Computer Vision and Pattern Recognition., (2018), pp. 586–595.

12. R. Girshick, J. Donahue, T. Darrell, J. Malik, Rich feature hierarchies for accurate object detection and semantic segmentation in Proceedings of the IEEE Conference on Computer Vision and Pattern Recognition (CVPR), Conference on Computer Vision and Pattern Recognition (CVPR)., (2014), pp. 580–587.

13. S. Kornblith, J. Shlens, Q. V. Le, Do Better ImageNet Models Transfer Better? in Proceedings of the IEEE Conference on Computer Vision and Pattern Recognition (CVPR), Conference on Computer Vision and Pattern Recognition (CVPR)., (2019), pp. 2661–2671.

14. M. Huh, P. Agrawal, A. A. Efros, What makes ImageNet good for transfer learning? in NIPS Workshop on Large Scale Computer Vision Systems, NIPS Workshop on Large Scale Computer Vision Systems., (2016), pp. 1–10.

15. L. Chang, B. Egger, T. Vetter, D. Y. Tsao, Explaining face representation in the primate brain using different computational models. bioRxiv (2021) https://doi.org/10.1101/2020.06.07.111930.

16. S. Grossman, et al., Convergent evolution of face spaces across human face-selective neuronal groups and deep convolutional networks. Nat. Commun. 10, 4934 (2019).

17. J. Yosinski, J. Clune, A. Nguyen, T. Fuchs, H. Lipson, “Understanding Neural Networks Through Deep Visualization” (2015).

18. S. Xu, Y. Zhang, Z. Zhen, J. Liu, “The face module emerged in a deep convolutional neural network selectively deprived of face experience” (2020).

19. S. Sutherland, B. Egger, J. Tenenbaum, “Building 3D Morphable Models from a Single Scan” (2020).

20. L. J. Powell, H. L. Kosakowski, R. Saxe, Social Origins of Cortical Face Areas. Trends Cogn. Sci. 22, 752–763 (2018).

21. A. Shoham, I. Grosbard, O. Patashnik, D. Cohen-Or, G. Yovel, Deep learning algorithms reveal a new visual-semantic representation of familiar faces in human perception and memory. Biorxiv, 2022.10.16.512398 (2022).

22. N. M. Blauch, M. Behrmann, D. C. Plaut, Computational insights into human perceptual expertise for familiar and unfamiliar face recognition. Cognition 208, 104341 (2020).

23. C. Rezlescu, A. Chapman, T. Susilo, A. Caramazza, Large inversion effects are not specific to faces and do not vary with object expertise. Preprint at PsyArXiv (2016) https://doi.org/10.31234/osf.io/xzbe5.

24. K. Simonyan, A. Zisserman, Very deep convolutional networks for large-scale image recognition in International Conference on Learning Representations, International Conference on Learning Representations., (2015), pp. 1–14.

25. M. Schrimpf, et al., “Brain-Score: Which Artificial Neural Network for Object Recognition is most Brain-Like?” (2018).

26. O. M. Parkhi, A. Vedaldi, A. Zisserman, Deep face recognition in Proceedings of the British Machine Vision Conference (BMVC), British Machine Vision Conference., (2015), p. 41.1–41.12.

27. A. Krizhevsky, I. Sutskever, G. E. Hinton, ImageNet Classification with Deep Convolutional Neural Networks in Adv. Neural Inf. Process. Syst., NIPS., (2012), pp. 1097–1105.

28. D. Han, J. Kim, J. Kim, Deep Pyramidal Residual Networks in Proceedings of the IEEE Conference on Computer Vision and Pattern Recognition (CVPR)), Conference on Computer Vision and Pattern Recognition (CVPR)., (2017), pp. 6307–6315.

29. K. He, X. Zhang, S. Ren, J. Sun, Deep Residual Learning for Image Recognition in Proceedings of the IEEE Conference on Computer Vision and Pattern Recognition (CVPR), Conference on Computer Vision and Pattern Recognition (CVPR)., (2016), pp. 770–778.

30. N. Abudarham, I. Grosbard, G. Yovel, Face recognition depends on specialized mechanisms tuned to view-invariant facial features: Insights from deep neural networks optimized for face or object recognition. Cogn. Sci. 45, e13031 (2021).

31. H. Katti, S. P. Arun, Are you from North or South India? A hard face-classification task reveals systematic representational differences between humans and machines. J Vision 19, 1 (2019).

32. C. M. Funke, et al., Five points to check when comparing visual perception in humans and machines. J. Vis. 21, 16–16 (2021).

33. N. Kriegeskorte, M. Mur, Inverse MDS: Inferring Dissimilarity Structure from Multiple Item Arrangements. Front. Psychol. 3, 245 (2012).

34. C. A. Meissner, J. C. Brigham, Thirty years of investigating the own-race bias in memory for faces: A meta-analytic review. Psychol. Public Policy Law 7, 3–35 (2001).

35. I. D. Raji, et al., Saving Face: Investigating the Ethical Concerns of Facial Recognition Auditing in AAAI/ACM Conference on AI, Ethics, and Society, (2020), pp. 145–151.

36. J. G. Cavazos, P. J. Phillips, C. D. Castillo, A. J. O’Toole, Accuracy comparison across face recognition algorithms: Where are we on measuring race bias? IEEE Trans. Biom. Behav. Identity Sci. 3, 101–111 (2021).

37. J. Tian, H. Xie, S. Hu, J. Liu, Multidimensional Face Representation in a Deep Convolutional Neural Network Reveals the Mechanism Underlying AI Racism. Front. Comput. Neurosci. 15, 620281 (2021).

38. I. Gauthier, C. A. Nelson, The development of face expertise. Curr. Opin. Neurobiol. 11, 219–224 (2001).

39. R. Robbins, E. McKone, No face-like processing for objects-of-expertise in three behavioural tasks. Cognition 103, 34–79 (2007).

40. A. Campbell, J. W. Tanaka, Inversion Impairs Expert Budgerigar Identity Recognition: A Face-Like Effect for a Nonface Object of Expertise. Perception 47, 647–659 (2018).

41. G. Schalk, et al., Facephenes and rainbows: Causal evidence for functional and anatomical specificity of face and color processing in the human brain. Proc. Natl. Acad. Sci. U.S.A. 114, 12285–12290 (2017).

42. K. Dobs, J. Martinez, A. J. E. Kell, N. Kanwisher, Brain-like functional specialization emerges spontaneously in deep neural networks. Sci. Adv. 8, eabl8913 (2022).

43. J. Buolamwini, T. Gebru, Gender Shades: Intersectional Accuracy Disparities in Commercial Gender Classification in Proceedings of Machine Learning Research, (2018), pp. 77–91.

44. G. Jacob, R. T. Pramod, H. Katti, S. P. Arun, Qualitative similarities and differences in visual object representations between brains and deep networks. Nat. Commun. 12, 1872 (2021).

45. G. Yovel, N. Abudarham, From concepts to percepts in human and machine face recognition: A reply to Blauch, Behrmann & Plaut. Cognition 208, 104424 (2020).

46. A. Farzmahdi, K. Rajaei, M. Ghodrati, R. Ebrahimpour, S.-M. Khaligh-Razavi, A specialized face-processing model inspired by the organization of monkey face patches explains several face-specific phenomena observed in humans. Sci. Rep. 6, 25025 (2016).

47. C. Zhuang, et al., Unsupervised neural network models of the ventral visual stream. Proc. Natl. Acad. Sci. U.S.A. 118, e2014196118 (2021).

48. N. Kanwisher, M. Khosla, K. Dobs, Using Artificial Neural Networks to Ask Why Questions of Minds and Brains. Trends in Neurosciences (2022).

49. L. Chang, B. Egger, T. Vetter, D. Y. Tsao, Explaining face representation in the primate brain using different computational models. Curr. Biol. (2021) https://doi.org/10.1016/j.cub.2021.04.014.

50. S. Anzellotti, A. Caramazza, Multimodal representations of person identity individuated with fMRI. Cortex 89, 85–97 (2017).

51. K. Dobs, J. Schultz, I. Bülthoff, J. L. Gardner, Task-dependent enhancement of facial expression and identity representations in human cortex. NeuroImage 172, 689–702 (2018).

52. J. S. Guntupalli, K. G. Wheeler, M. I. Gobbini, Disentangling the Representation of Identity from Head View Along the Human Face Processing Pathway. Cereb. Cortex 27, 46–53 (2016).

53. N. Kriegeskorte, E. Formisano, B. Sorger, R. Goebel, Individual faces elicit distinct response patterns in human anterior temporal cortex. Proc. Natl. Acad. Sci. U.S.A. 104, 20600–20605 (2007).

54. S. M. Landi, W. A. Freiwald, Two areas for familiar face recognition in the primate brain. Science 357, 591–595 (2017).

55. J. Dubois, A. O. de Berker, D. Y. Tsao, Single-Unit Recordings in the Macaque Face Patch System Reveal Limitations of fMRI MVPA. J. Neurosci. 35, 2791–2802 (2015).

56. T. C. Kietzmann, et al., Recurrence is required to capture the representational dynamics of the human visual system. Proc. Natl. Acad. Sci. U.S.A. 116, 21854– 21863 (2019).

57. K. Kar, J. Kubilius, K. Schmidt, E. B. Issa, J. J. DiCarlo, Evidence that recurrent circuits are critical to the ventral stream’s execution of core object recognition behavior. Nat. Neurosci. 22, 974–983 (2019).

58. I. Yildirim, M. Belledonne, W. Freiwald, J. Tenenbaum, Efficient inverse graphics in biological face processing. Sci. Adv. 6, eaax5979 (2020).

59. K. M. Jozwik, et al., Face dissimilarity judgments are predicted by representational distance in morphable and image-computable models. Proc National Acad Sci 119, e2115047119 (2022).

60. C. Daube, et al., Grounding deep neural network predictions of human categorization behavior in understandable functional features: The case of face identity. Patterns 2, 100348 (2021).

61. A. Francl, J. H. McDermott, Deep neural network models of sound localization reveal how perception is adapted to real-world environments. Nat. Hum. Behav. 6, 111–133 (2022).

62. M. R. Saddler, R. Gonzalez, J. H. McDermott, Deep neural network models reveal interplay of peripheral coding and stimulus statistics in pitch perception. Nat. Commun. 12, 7278 (2021).

63. M. N. Hebart, C. Y. Zheng, F. Pereira, C. I. Baker, Revealing the multidimensional mental representations of natural objects underlying human similarity judgements. Nat. Hum. Behav. 4, 1173–1185 (2020).

64. G. B. Huang, M. Mattar, T. Berg, E. Learned-Miller, Labeled Faces in the Wild: A Database for Studying Face Recognition in Unconstrained Environments in Workshop on Faces in “Real-Life” Images: Detection, Alignment, and Recognition, (2008), pp. 1–11.

65. J. Deng, et al., ImageNet: A large-scale hierarchical image database in Proceedings of the IEEE Conference on Computer Vision and Pattern Recognition, Conference on Computer Vision and Pattern Recognition (CVPR)., (2009), pp. 248– 255.

66. R. Jenkins, A. J. Dowsett, A. M. Burton, How many faces do people know? Cereb. Cortex 285, 20181319–6 (2018).

67. K. Dobs, L. Isik, D. Pantazis, N. Kanwisher, How face perception unfolds over time. Nat. Commun. 10, 1258 (2019).

68. H. Nili, et al., A Toolbox for Representational Similarity Analysis. PLoS Comp. Biol. 10, e1003553–11 (2014).

69. T. Karras, S. Laine, T. Aila, A Style-Based Generator Architecture for Generative Adversarial Networks in Proceedings of the IEEE Conference on Computer Vision and Pattern Recognition (CVPR), (2019), pp. 4401–4410.

70. Z. Xiong, et al., An Asian Face Dataset and How Race Influences Face Recognition in Pacific Rim Conference on Multimedia, Pacific Rim Conference on Multimedia., (2018), pp. 372–383.

71. L. Yang, P. Luo, C. C. Loy, X. Tang, A Large-Scale Car Dataset for Fine-Grained Categorization and Verification in Proceedings of the IEEE Conference on Computer Vision and Pattern Recognition, (2015), pp. 3973–3981.

