## Supplementary Information for "Behavioral signatures of face perception emerge in deep neural networks optimized for face recognition"

Dobs et al.

**Table of contents**

**Supplementary Note 1:** Layer-wise analysis of face recognition performance, other-race and face inversion effect in VGG16
**Supplementary Note 2:** Replication of results for face recognition performance, other-race and face inversion effect in Alexnet and ResNet
**Supplementary Note 3:** Analysis of errors made by humans and by different CNNs
**Supplementary Note 4:** RDMs of representational similarity analyses in VGG16
**Supplementary Note 5:** Layer-wise representational similarity analysis in VGG16
**Supplementary Note 6:** Representational similarity analysis in Alexnet and ResNet
**Table S1:** Overview of experiments, participants and datasets
**Table S2:** Overview of experimental datasets
**Table S3:** Overview of CNN experiments
**Table S4:** Overview of trained and untrained CNNs

**Supplementary Note 1: Layer-wise analysis of face recognition performance, other-race and face inversion effect in VGG16**

**Methods.** To compare the face recognition performance between humans and CNNs for each layer, we extracted activation from each layer of three VGG16 networks (Face-ID CNN; Obj-Cat CNN and Untrained CNN) and performed the same analysis. Specifically, we extracted the activation from each convolutional and fully-connected layer after the relu operation. When the layer was followed by a pooling layer, we extracted the activation from the pooling layer. For each model and layer, we computed the correlation distance between the activation patterns of each pair of images (Fig. 1B, right panel). The network's choice was determined by which of the two matching images had an activation pattern that was closest to the target image. We performed this analysis for the white female datasets upright and inverted, and for the unfamiliar white and Asian female datasets.

**Results.** For the white female dataset, we found that the face-trained CNN began to outperform the object-trained CNN from the last convolutional layer onwards (Supplementary Fig. 1; p=0, bootstrap test). The performance of the untrained and the object-trained CNNs did not vary much across layers. These results show that the late stages of the face-trained network outperform the object-trained network and approach human-level face recognition performance.

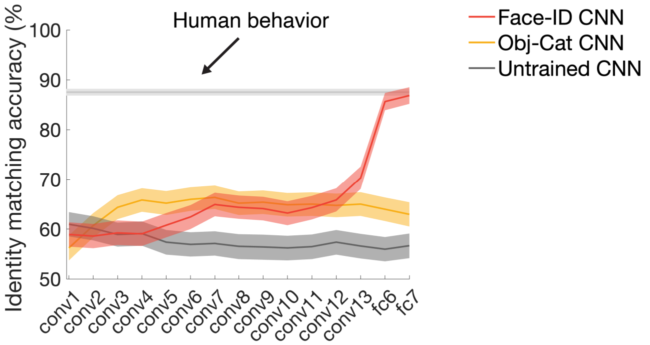


**Supplementary Figure 1 | Late layers of face-trained CNNs outperform object-trained and untrained CNNs and achieve humanlike performance. (a)** Human performance (n=1,552) was 87.53% (gray horizontal line; chancel level was 50%). The face-trained CNN (red) outperformed the object-trained CNN (yellow) from the last convolutional layer (conv13) onwards (p=0, bootstrap test). The face-trained CNN further reached human performance in the penultimate fully-connected layer (fc7; p>0.4, bootstrap test). Networks trained on object categorization (yellow) performed better than untrained CNNs (gray) across all layers, but did not reach human-level recognition performance. Error bars denote bootstrapped 95% CI.

To measure the other-race effect in each layer, we compared the layer-wise accuracy on the target-matching task on the non-famous white female dataset to the accuracy on the Asian female dataset (Supplementary Fig. 2). The performance of the white face-trained network on white female dataset was significantly higher than for the Asian female dataset from the last convolutional layer onwards (p=0, bootstrap test) and vice versa for the Asian face trained CNN. Neither the object-trained nor the untrained CNN showed a significant different between the two datasets (all p>0.2, bootstrap test).

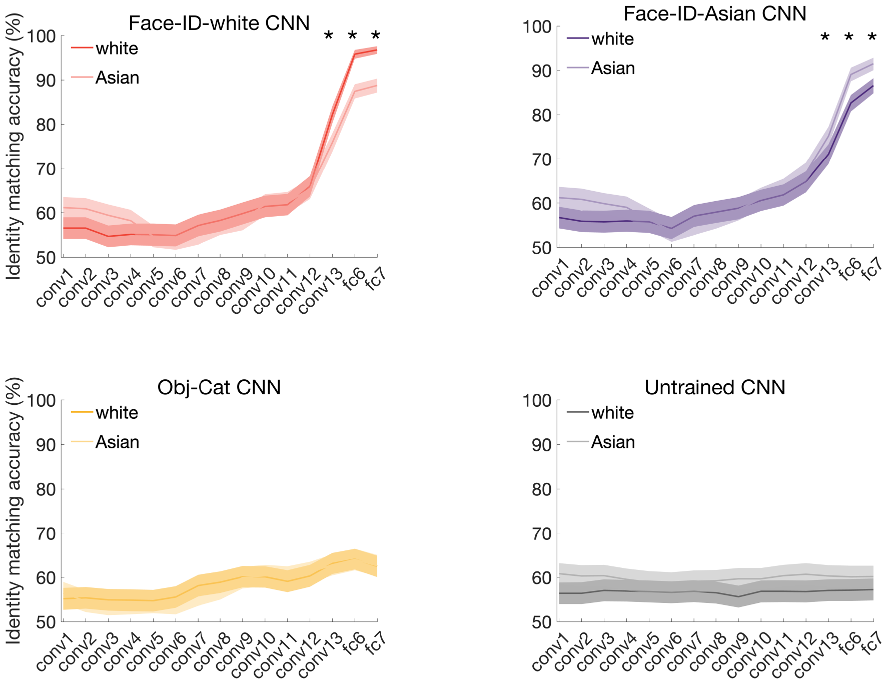


**Supplementary Figure 2 | Late layers of the face-trained CNNs show an other-race effect.** For the face-trained CNN (Face-white CNN in red; all Asian faces were removed from the training), the performance on the white female dataset was significantly higher than on the Asian female dataset from the last convolutional layer onwards. The opposite was true for the CNN trained on Asian identities (Face-Asian CNN in purple). Neither the object-trained (yellow) nor the untrained (gray) CNN showed a significant difference between both datasets at any layer. Shaded areas denote bootstrapped 95% CI. Stars indicate significant differences (bootstrap test, p<1e-5).

We found similar results when comparing the performance on the upright and inverted white female dataset (Supplementary Fig. 3). Only the face-trained, but not the object-trained or untrained, CNN showed a significant face inversion effect, i.e., improved performance for upright compared to inverted images, from the last convolutional layer onwards (p<0.002, bootstrap test).

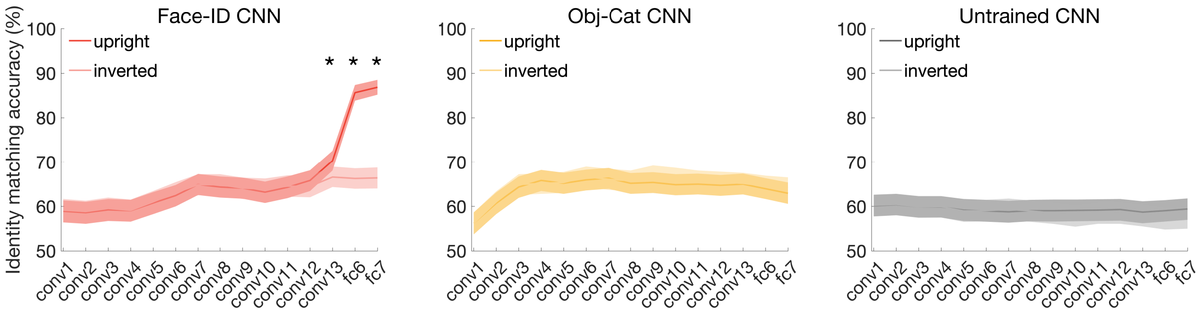


**Supplementary Figure 3 | Late layers of the face-trained CNN show a face inversion effect.** The pattern of differences was very similar to the other-race effect. Only the face-trained CNN showed a significant difference in performance between upright and inverted face stimuli starting at the last convolutional layer. Shaded areas denote bootstrapped 95% CI. Stars indicate significant differences (bootstrap test, p<1e-4).

### Supplementary Note 2: Replication of results for face recognition performance, other-race and face inversion effect in Alexnet and ResNet

**Methods.** To test whether the main results in the paper generalize to other architectures, we compared face recognition performance of humans to Alexnet and ResNet-50. For each of these architectures, we trained three different networks: i) one trained on face discrimination, ii) one trained on object categorization, and iii) one untrained. Note that for computational efficiency, we restricted this analysis to the face-trained, object-trained and untrained CNNs (referred to as Face CNN, Object CNN, Untrained CNN, respectively). We used the same training stimuli, parameters and procedure to train the CNNs on face or object categorization. For both architectures, we extracted the activation from the last relu (Alexnet) or the last pooling (ResNet) layer preceding the classification layer. We performed this analysis for the white female identities upright and inverted, and for the non-famous white and Asian female identities, respectively.

**Results.** The pattern of results on the white female dataset obtained for VGG16 were generally replicated with Alexnet and ResNet (Supplementary Fig. 4). We found that the face-trained CNN (red) outperformed the object-trained CNN for both architectures ($p=0$, bootstrap test), while the object-trained CNN outperformed the untrained CNN ($p=0$, bootstrap test). However, neither Alexnet nor ResNet reached human performance ($p=0$, bootstrap test).

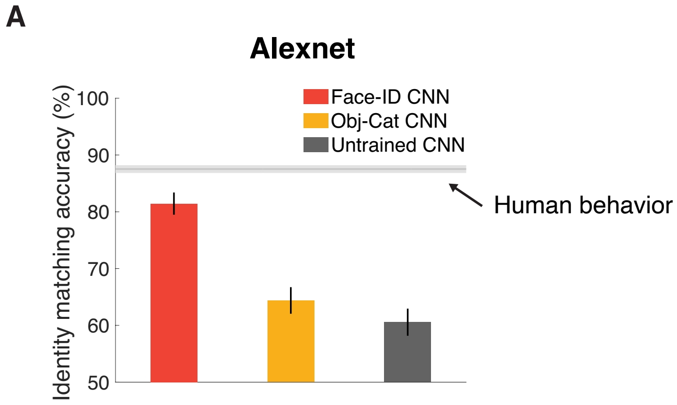


**Supplementary Figure 4 | Pattern of face recognition results replicated in Alexnet and ResNet.** Error bars denote bootstrapped 95% CI.

To investigate the face inversion effect in Alexnet and ResNet, we ran the same analysis on the inverted face dataset (Supplementary Fig. 5). As we found for VGG16, the performance of the face-trained Alexnet and ResNet on upright stimuli was significantly higher than for inverted stimuli (p=0, bootstrap test). In both architectures, neither the object-trained nor the untrained CNN showed a significant difference between the two datasets (all p>0.2, bootstrap test). These results show that both architectures, if trained on (upright) faces, also show a face inversion effect.

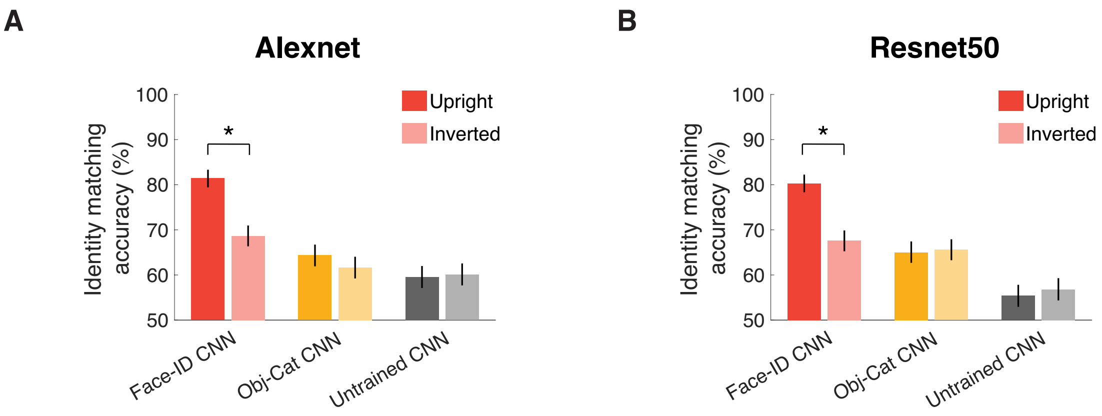


**Supplementary Figure 5 | Face inversion effect in Alexnet and ResNet.** (a) For Alexnet trained on (upright) faces (red), the performance on the upright female dataset was significantly higher than on the inverted female dataset in the penultimate fully-connected layer. Neither the object-trained (yellow) nor the untrained (gray) Alexnet showed a significant difference between the two datasets. (b) The pattern of differences was very similar for ResNet and replicated the results for Alexnet and VGG16: Only the face-trained CNN showed a significant difference in performance between upright and inverted face stimuli in the penultimate fully-connected layer. Error bars denote bootstrapped 95% CI. Asterisks indicate significant differences (bootstrap test, p=0).

We found very similar results when comparing the performance on the non-famous white female and Asian female dataset (Supplementary Fig. 6). Only the face-trained, but not the object-trained or untrained, architectures showed a significant other-race effect, that is improved performance for white compared to Asian face images in the penultimate fully-connected layer (p=0, bootstrap test).

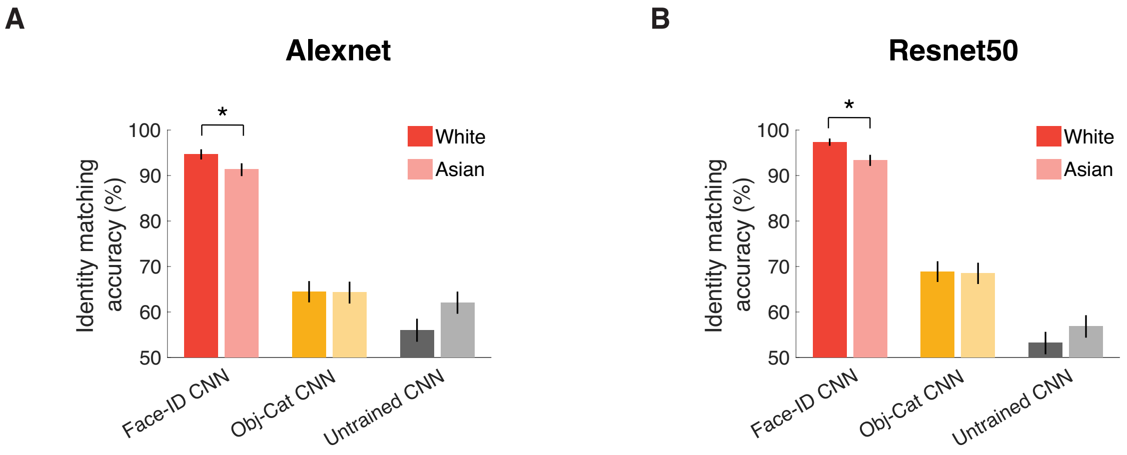


**Supplementary Figure 6 | Other-race effect in Alexnet and ResNet. (a)** For Alexnet trained on (predominantly white) faces (red), the performance on the white female dataset was significantly higher than on the Asian female dataset in the penultimate fully-connected layer. Neither the object-trained (yellow) nor the untrained (gray) Alexnet showed a significant difference between the two datasets. **(b)** The pattern of differences for the ResNet architecture was very similar to Alexnet and replicated the results for VGG16: Only the face-trained CNN showed a significant difference in performance between white and Asian face stimuli in the penultimate fully-connected layer. Error bars denote bootstrapped 95% CI. Asterisks indicate significant differences (bootstrap test, p=0).

**Supplementary Note 3: Analysis of errors made by humans and by different CNNs**

**Methods.** The analyses so far show that CNNs trained on face recognition achieve accuracy levels similar to humans when tested on the same task. But do they achieve this in the same way? To address this question, we asked whether humans and CNNs find the same triplets (i.e., a specific combination of target and two matching images) difficult. To test whether triplets that the CNNs did not perform correctly were also harder for human participants, we separately analyzed the triplets for which the CNNs were correct versus incorrect.

**Results.** For the face-identity trained CNN, we indeed found that human performance was significantly better on triplets in which the CNN was correct (human performance 88.2%; 1355 triplets) than on triplets for which it was incorrect (human performance: 83.3%; 205 triplets; p=0, bootstrap test). Moreover, this difference in performance (4.9%) was significantly smaller for the CNN trained on object categorization (difference: 1.5%; p=0.02, bootstrap test) and the untrained CNN (difference: 1.6%; p=0.02, bootstrap test). Here, humans performed significantly but only slightly worse on triplets the CNNs performed incorrectly (human performance: 86.6% for both Obj-Cat and untrained CNN) than on triplets the CNNs performed correctly (human performance: 88.1% for Obj-Cat CNN; p=0.03, bootstrap test; 88.2% for Untrained CNN; p=0.02, bootstrap test). This finding suggests that the face-identity trained CNN not only achieves a similar recognition accuracy to humans, but also shows similar errors to humans.

Would a network trained on face detection better match human face behavior? We indeed found a significant difference of 3.2% in human performance on triplets in which the CNN trained on object and face categorization (Obj-Face-Cat CNN) was correct vs. incorrect (human performance: 88.6% (1047 triplets) vs. 85.4% (513 triplets); p=0, bootstrap test). The size of this difference was not significantly different from those found for any of the other CNNs (Face-ID CNN: p=0.2; Obj-Cat and Untrained CNN: p=0.1; bootstrap tests).

**Supplementary Note 4: RDMs of representational similarity analyses in VGG16**

**Methods.** We used representational similarity analysis to compare the behavioral representational dissimilarity matrices (RDMs) obtained from the multi-arrangement task (Experiment 2; Fig. 2B) and the similarity-matching task (Experiment 3) to the RDMs of all four VGG16 models. Here, we plot the behavioral RDMs along with the RDMs obtained from the four VGG16 models for visual comparison.

**Results.** For the multi-arrangement task (Experiment 2; Supplementary Fig. 7), the face-trained CNN (Face-ID CNN) best mirrors the structure of the behavioral RDMs. This similarity goes beyond the coarse distinctions between male and female and old and young faces. For example, within the old female faces, the Face-ID CNN also shows a high similarity between the third and the fourth identity. Note that while the object-trained and object-and-face-categorization trained CNNs also show some of the coarse categories (e.g., older male faces in the bottom right are highly similar to each other but distinct from young female faces), they do not capture all aspects of the fine-grained structure. For example, older female faces are more similar to young male faces (blue colors in the bottom left quadrant) in these CNNs than in human behavior and the Face-ID CNN (yellow colors in the bottom left quadrant). Interestingly, even the untrained CNN shows a trend for some of the aspects in the behavioral RDM, such as a high similarity between older female identities.

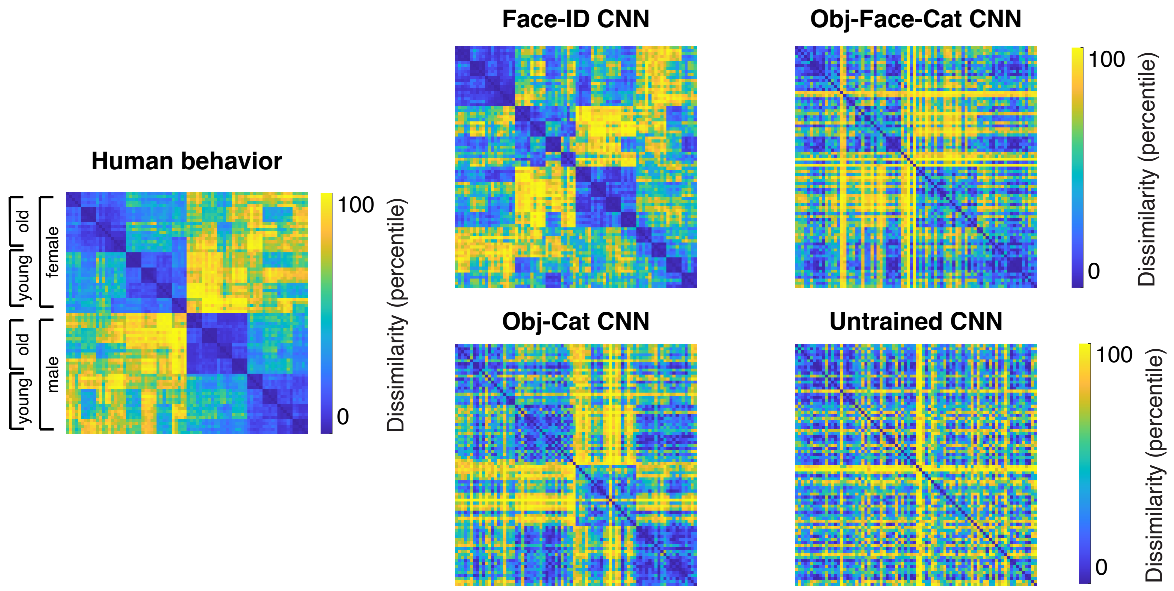


**Supplementary Figure 7 | Face-trained CNN best matches human perceptual similarity in a multi-arrangement task (Experiment 2).** The stimuli used in the multi-arrangement task consisted of 5 stimuli for each of 16 identities for which half of them were old versus young and female versus male, respectively. In the RDM, the stimuli are arranged by identity (5 images per identity), age (young versus old) and gender (female versus male). The Face-ID CNN best matches the behavioral RDM. The Obj-Face-Cat and Obj-Cat CNN show some coarse aspects but do not match the fine-grained details of the behavioral RDM.

For the similarity-matching task (Experiment 3; Supplementary Fig. 8), all 60 identities were young and male and we used one image per identity. As can be seen from the behavioral RDM, there is no clear structure in the RDM. As for the multi-arrangement ask, the RDM of the face-trained CNN appears most similar to the behavioral RDM. In particular, the other three CNNs show strong similarities for certain images with all other images (as can evident by blue lines in the matrices), which are not visible in the behavioral or the RDM of the Face-ID CNN.

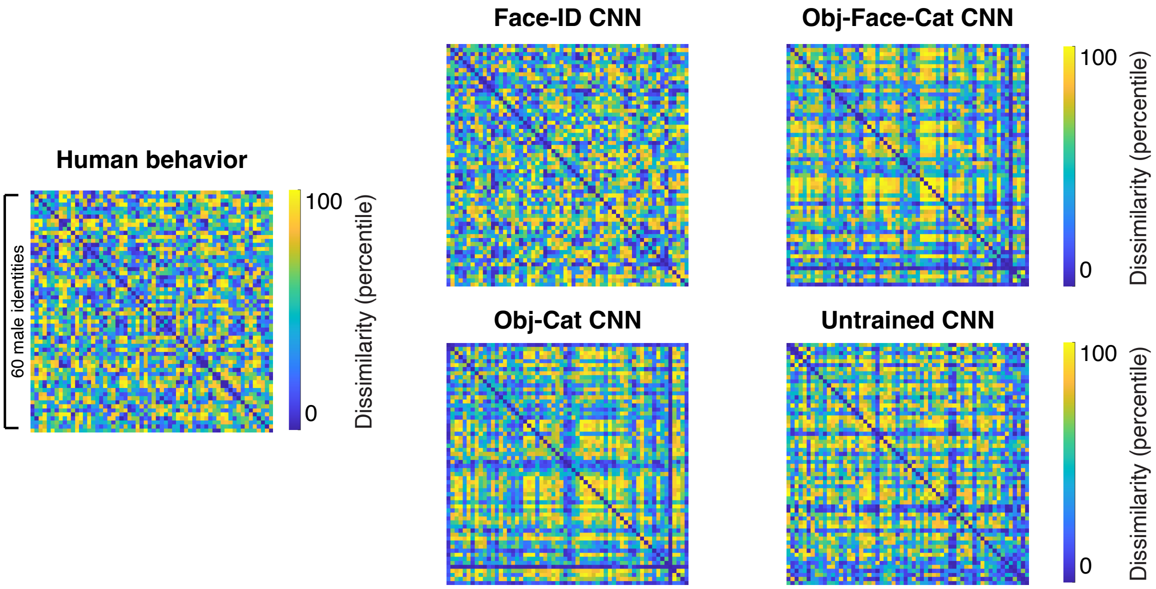


**Supplementary Figure 8 | Face-trained CNN best matches human perceptual similarity in a similarity-matching task (Experiment 2).** The stimuli in the similarity-matching task consisted of 60 young, male identities (one image per identity). The Face-ID CNN best matches the behavioral RDM. The Obj-Face-Cat and Obj-Cat CNN show specific structures (e.g., strong similarity of a specific image with all other images) that are neither visible in the behavioral RDM nor the face-trained CNN.

**Supplementary Note 5: Layer-wise representational similarity analysis in VGG16**

**Methods.** We used representational similarity analysis to compare the behavioral representational dissimilarity matrices (RDMs) obtained from the multi-arrangement task (Experiment 2; Fig. 2B) and the similarity-matching task (Experiment 3; Fig. 2C) to the layer-wise RDMs of the VGG16 models trained on face identification and object categorization and the untrained VGG16 model. Specifically, to obtain layer-wise RDMs for each model, we computed the distance (i.e., 1 – Pearson's *r*) between the activation patterns extracted from each layer for the same stimuli.

**Results.** In the multi-arrangement task (Experiment 2; Supplementary Fig. 9a), we find that correlations between the face-trained CNN (red) and human behavior increased with progressive layers in the network from the first convolutional layer (Spearman's *r*: 0.05), to mid-level convolutional layers (e.g., Conv8: Spearman's *r*: 0.16) to the last convolutional layer (Spearman's *r*: 0.34), the latter even reaching noise ceiling (i.e., the maximum correlation possible given the consistency across subjects; light-gray vertical bar). In contrast, the object-trained CNN (yellow) represented faces less similarly to humans (max. Spearman's *r*: 0.19), with correlations increasing slightly after the first 4-5 layers, reaching its maximum in the penultimate fully-connected layer. The representational dissimilarities of the untrained CNN (dark gray) showed a low correlation with human behavior across all layers (max. Spearman's *r*: 0.04). Thus, the later stages of processing in face-trained, but not object-trained or untrained, CNNs match human behavior well, suggesting that faces are similarly represented in human behavior and late stages of face-trained CNNs.

We find a very similar pattern for the similarity-matching task (Experiment 3; Supplementary Fig. 9b).

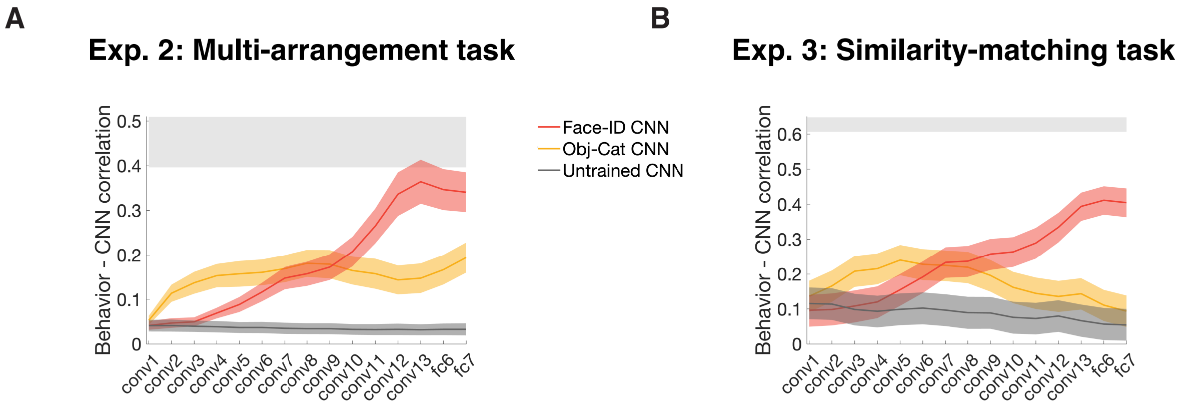


**Supplementary Figure 9 | Late layers of face-trained but not object-trained or untrained CNNs match human face behavior. (a)** We performed RSA on all layers of the three VGG16 models and human behavioral similarities from the multi-arrangement task (Experiment 2). Late layers of the Face CNN (red) matched human behavioral representational similarity best and reached the noise ceiling (light gray bar). Neither the untrained CNN (Untrained CNN, dark gray) nor the object-trained CNN (Object CNN, yellow) matched human representational similarities. Shaded areas represent bootstrapped SEMs across subjects. **(b)** The results in (a) were replicated on the similarity-matching task (Experiment 3) using on a distinct dataset of 60 unfamiliar male identities (one image each). Late layers of the Face CNN (red) matched human behavioral representational similarity best, far outperforming the untrained CNN (gray) and the object-trained CNN (yellow). Shaded areas represent bootstrapped SEMs across triplets.

**Supplementary Note 6: Representational similarity analysis in Alexnet and ResNet**

**Methods.** To test whether these results would generalize to other architectures, we compared the behavioral RDMs obtained from the multi-arrangement task (Experiment 12 Fig. 2A) and the similarity-matching task (Experiment 3) to the CNN RDMs of all three Alexnet and ResNet-50 models. Specifically, to obtain RDMs for each model, we computed the distance (i.e., 1 – Pearson's *r*) between the activation patterns extracted from the penultimate layer for the same stimuli.

**Results.** For the multi-arrangement task (Supplementary Fig. 10a), the correlations between the face-trained Alexnet (red) and human behavior were close to the noise ceiling (Spearman's *r*: 0.32, close to noise ceiling). In contrast, the object-trained CNN (yellow) represented faces less similarly to humans (Spearman's *r*: 0.18). The representational dissimilarities of the untrained CNN (dark gray) showed a low correlation with human behavior (Spearman's *r*: 0.04). We replicated this pattern of results for Alexnet in the similarity-matching task (Supplementary Fig. 10b ). Thus, processing in face-trained, but not object-trained or untrained, Alexnet models match human behavior well.

ResNet trained on faces, objects and untrained showed a very similar pattern. In the multi-arrangement task (Supplementary Fig. 10a) the face-trained CNN (red) even reached the noise ceiling (Spearman's *r*: .36), while the object-trained (Spearman's *r*: .19) and the untrained (Spearman's *r*: .03) CNN achieved much lower correlations with human similarity representations. We again replicated this pattern of results for ResNet in the similarity-matching task (Supplementary Fig 10b).

Taken together, these findings suggest that faces are similarly represented in human behavior and face-trained feed-forward CNNs, irrespective of architecture.

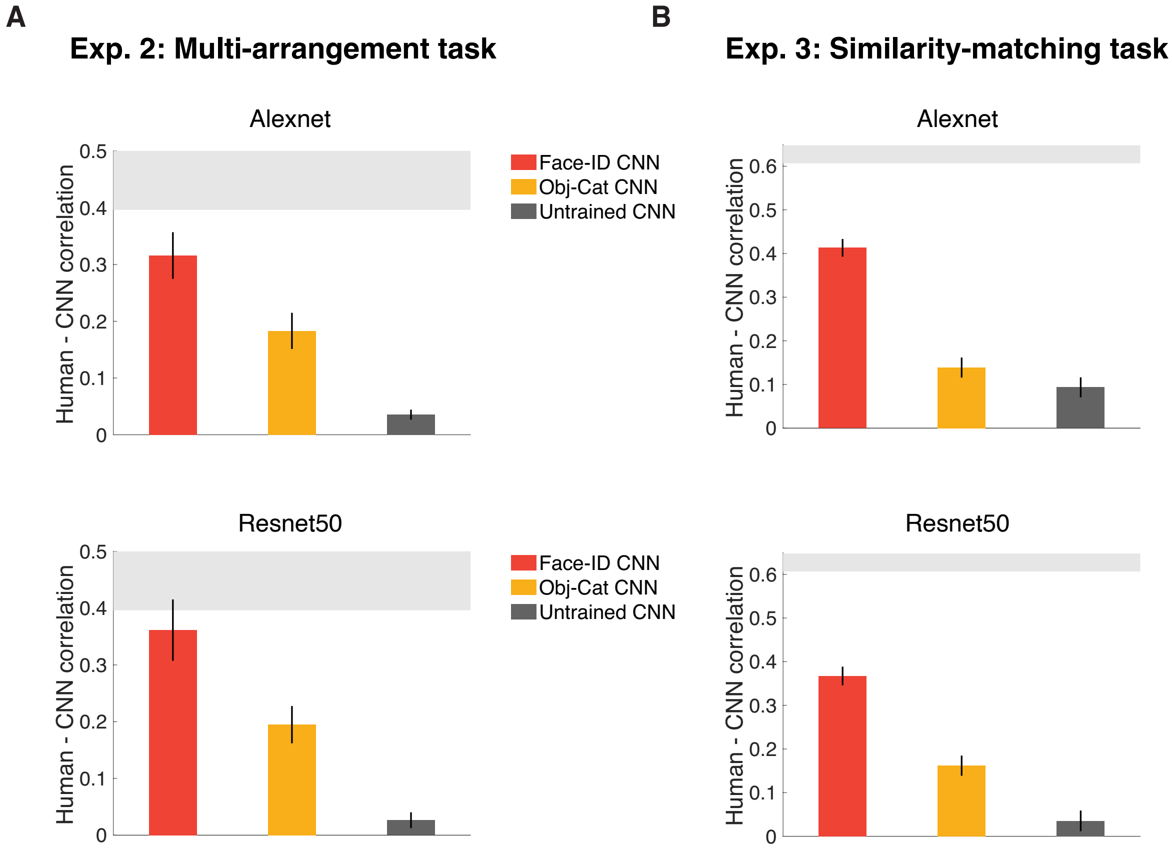


**Supplementary Figure 10 | Face-trained but not object-trained or untrained Alexnet and ResNet-50 architectures approach human face behavior. (a)** We performed RSA by measuring the similarity between human behavioral similarities from a multi-arrangement task (Experiment 2) and layer-wise RDMs obtained from the three Alexnet and three ResNet-50 models. For both architectures, the face-trained models (red) matched human behavioral representational similarity best and approached noise ceiling (light gray bar). Neither the untrained CNN (dark gray) nor the object-trained CNN (yellow) matched human representational similarities. Error bars represent bootstrapped SEMs across subjects. **(b)** The results of in (a) were replicated in a similarity-matching task on Amazon Mechanical Turk (Experiment 3) based on a distinct dataset of 60 unfamiliar male identities (one image each). For both architectures, the Face CNN (red) matched human behavioral representational similarity best, far outperforming the untrained CNN (gray) and the object-trained CNN (yellow). Error bars represent bootstrapped SEMs across stimuli.

**Table S1.** Overview of experiments, participants and datasets

| Human Experiment | Human participants (n) | Testing platform | Task | Stimuli | Figure |
|---|---|---|---|---|---|
| Exp. 1: Face recognition - upright | 1,532 | Amazon Mechanical Turk | Target-matching task | Set A | Fig. 1C |
| Exp. 2: Perceptual similarity | 14 | Meadows | Multi-arrangement task | Set B | Fig. 2B |
| Exp. 3: Perceptual similarity | 668 | Amazon Mechanical Turk | Similarity-matching task | Set C | Fig. 2C |
| Exp. 4A: Other-race effect - white participants | 269 | Amazon Mechanical Turk | Target-matching task | Set D | Fig. 3A |
| Exp. 4B: Other-race effect - Asian participants | 102 | Clickworker + Meadows | Target-matching task | Set D | Fig. 3A |
| Exp. 5: Face recognition - inverted | 1,219 | Amazon Mechanical Turk | Target-matching task | Set A inverted | Fig. 3B |

**Table S2.** Overview of experimental datasets

| **Test Stimulus Set Name** | **Description** | **Link** |
|---|---|---|
| Set A | 200 face images (5 images of each of 40 celebrities) | https://osf.io/dbks3/ |
| Set B | 80 face images (5 images of each of 16 identities) | https://osf.io/gk6f5/ |
| Set C | 60 face images (1 image of each of 60 identities) | https://osf.io/dbks3/ |
| Set D | 400 face images (5 images of each of 40 white and 40 Asian identities) | https://osf.io/dbks3/ |
| Set E | 1000 face images (10 images of each 100 identities) | https://osf.io/dbks3/ |
| Set F | 1000 car images (10 images of each 100 car model/makes) | https://osf.io/dbks3/ |

**Table S3.** Overview of CNN experiments

| CNN Experiment | CNNs | Analysis Method | Stimuli | Figure |
|---|---|---|---|---|
| Face recognition – upright (Exp. 1) | CNN 1-4 | Target-matching task | Set A | Fig. 1C |
| Perceptual similarity (Exp. 2) | CNN 1-4 | RSA | Set B | Fig. 2B |
| Perceptual similarity (Exp. 3) | CNN 1-4 | RSA | Set C | Fig. 2C |
| Other-race effect (Exp. 4) | CNN 3-8 | Target-matching task | Set D | Fig. 3A |
| Face recognition – inverted (Exp. 5) | CNN 1-4 | Target-matching task | Set A inverted | Fig. 3B |
| Inverted face inversion effect | CNN 1, 9 | SVM decoding | Set E | Fig. 4A |
| Car inversion effect | CNN 1, 3, 4, 10 | SVM decoding | Set F | Fig. 4B |

**Table S4.** Overview of trained and untrained CNNs

| CNN # | CNN Name | Training Set | Link |
|---|---|---|---|
| CNN 1 | Face-ID CNN | 1,714 VGGFace2 classes | https://github.com/ox-vgg/vgg_face2 |
| CNN 2 | Obj-Face-Cat CNN | 423 ImageNet classes + 1,714 VGGFace2 classes (assigned to one output class) | |
| CNN 3 | Obj-Cat CNN | 423 ImageNet classes | https://www.image-net.org/challenges/LSVRC/2012/index.php |
| CNN 4 | Untrained CNN | None | |
| CNN 5 | Face-ID-white CNN | 1,654 VGGFace2 classes (white only) | |
| CNN 6 | Face-ID-Asian CNN | 1,654 Asian Face Dataset classes | https://github.com/X-zhangyang/Asian-Face-Image-Dataset-AFD-dataset |
| CNN 7 | Obj-Face-Cat-white CNN | 423 ImageNet classes + 1,654 VGGFace2 classes (white only; assigned to one output class) | |
| CNN 8 | Obj-Face-Cat-Asian CNN | 423 ImageNet classes + 1,654 Asian Face Dataset classes (assigned to one output class) | |
| CNN 9 | Face-ID-inv CNN | 1,714 VGGFace2 classes (inverted) | |
| CNN 10 | Car CNN | 1,109 combined CompCars dataset classes | http://mmlab.ie.cuhk.edu.hk/datasets/comp_cars/ |
